# Single-Cell Profiling of the Antigen-Specific Response to BNT162b2 SARS-CoV-2 RNA Vaccine

**DOI:** 10.1101/2021.07.28.453981

**Authors:** Kevin J. Kramer, Erin M. Wilfong, Kelsey Voss, Sierra M. Barone, Andrea R. Shiakolas, Nagarajan Raju, Caroline E. Roe, Naveenchandra Suryadevara, Lauren Walker, Steven C. Wall, Ariana Paulo, Samuel Schaefer, Debolanle Dahunsi, Camille S. Westlake, James E. Crowe, Robert H. Carnahan, Jeffrey C. Rathmell, Rachel H. Bonami, Ivelin S. Georgiev, Jonathan M. Irish

## Abstract

RNA-based vaccines against SARS-CoV-2 are critical to limiting COVID-19 severity and spread. Cellular mechanisms driving antigen-specific responses to these vaccines, however, remain uncertain. We used single-cell technologies to identify and characterized antigen-specific cells and antibody responses to the RNA vaccine BNT162b2 in longitudinal samples from a cohort of healthy donors. Mass cytometry and machine learning pinpointed a novel expanding, population of antigen-specific non-canonical memory CD4^+^ and CD8^+^ T cells. B cell sequencing suggested progression from IgM, with apparent cross-reactivity to endemic coronaviruses, to SARS-CoV-2-specific IgA and IgG memory B cells and plasmablasts. Responding lymphocyte populations correlated with eventual SARS-CoV-2 IgG and a donor lacking these cell populations failed to sustain SARS-CoV-2-specific antibodies and experienced breakthrough infection. These integrated proteomic and genomic platforms reveal an antigen-specific cellular basis of RNA vaccine-based immunity.

**ONE SENTENCE SUMMARY:** Single-cell profiling reveals the cellular basis of the antigen-specific response to the BNT162b2 SARS-CoV-2 RNA vaccine.

## INTRODUCTION

In December 2019, a novel coronavirus strain designated severe respiratory distress syndrome coronavirus 2 (SARS-CoV-2) was identified in Wuhan, China. A global pandemic ensued that has resulted in approximately 200 million cases and over 4 million deaths of coronavirus disease 2019 (COVID-19) to date (*1*). B cells, T cells, and other leukocytes undergo significant shifts upon SARS-CoV-2 infection that may contribute to anti-viral immunity and protective antibodies (*2-6*). The development of viral neutralizing antibodies following infection has been associated with Th1-like CXCR3+HLADR+PD1+ CD8 and CD4 cells and circulating CXCR3+ CD4 T follicular helper (cTfh) cells and CD4+ CD38+ HLA-DR+ T cell abundance (*3, 4*). While therapies, such as dexamethasone (*7*), baracitinib (*8*), tocilizumab (*9*), and neutralizing monoclonal antibodies (*10, 11*) have emerged as treatments for severe COVID-19 disease, preventive measures to develop coronavirus immunity on a population-scale are of upmost importance. To address this need, vaccines formulated with the pre-fusion stabilized SARS-CoV-2 Spike (S) protein were developed to induce protection from COVID-19 infection or development of severe disease (*12-16*). Globally, nearly 4 billion doses of various COVID-19 vaccines have been administered (*1*).

Messenger RNA (mRNA)-based vaccines represent a promising new class of vaccines that offer protection from COVID-19 as well as potentially a wide range of emerging infectious diseases (*17, 18*). These vaccines introduce the minimal genetic information to express viral antigens of interest (*17*) and mimic natural infection of RNA viruses, such as SARS-CoV-2 (*19*). Several groups have explored the immunologic response to SARS-CoV-2 mRNA vaccines using both systems (*20*) and T-cell centric approaches (*21-23*). From these studies, elevated myeloid and T cell responses have been identified following vaccination and which corresponded to serologic antibody responses. However, antigen-specific cells that respond to vaccination and mechanisms of mRNA-based formulations remain poorly understood. It is also not well established how pre-existing immunity to endemic coronaviruses impacts the B cell and antibody response against SARS-CoV-2 vaccination and how the antibody repertoire may evolve over time

A major challenge to studies of immune responses to emerging diseases is reliable identification of antigen specific cells. While MHC tetramers and other tracking agents can identify pre-determined subsets of antigen-specific cells, a systemic and unbiased view of SARS-CoV-2 antigen responsive cells is needed. Single cell machine learning analysis tools including the Tracking Responders EXpanding (T-REX) algorithm (*24*) and Linking B cell Receptor to Antigen specificity through sequencing (LIBRA-seq) (*25*) combined with whole transcriptome RNA-seq may address this need to reveal immune cells reacting to infection or vaccination. These single-cell approaches identify rare cells that specifically expand following vaccination or infection that can be overlooked when analyzing cellular populations in bulk. Proteomic signatures identified by T-REX can be combined with Marker Enrichment Modeling (MEM) (*26*) to develop strategies to physically isolate the cell subset using fluorescence activated cell sorting (FACS) enabling *in vitro* validation. LIBRA-seq identifies antigen-binding B cells and their associated B-cell receptor (BCR) sequences, which can be recombinantly expressed as antibody and characterized *in vitro* together with single cell RNAseq to identify and characterize B cell subsets based on transcriptional profiles (*25*). Adaptation and evolution of antibody isotypes and antigen-binding properties provide further insight into the development of B cell responses and antigen-specificity.

To better understand the basis of antigen-specific cells and antibody response in SARS-CoV-2 vaccine-induced immunity, we tracked the development of antigen-specific T and B cells in longitudinal samples collected from healthy donor recipients of the BNT162b2 RNA-based SARS-CoV-2 vaccine, including a donor with a breakthrough SARS-CoV-2 infection. T-REX identified novel expanding and metabolically active S protein-specific, non-canonical memory CD4 and CD8 T cell populations following vaccination that were confirmed as antigen specific. In parallel, coronavirus S protein-binding B cells were characterized with single-cell LIBRA-seq and RNA-seq to establish the evolution of cross-reactive to antigen specific B cells with public antibody sequences over time. These antigen-specific T and B cells correlated and associated with a long-lasting IgG response that was lacking in a donor who subsequently experienced a breakthrough infection. These cell and antibody associations may drive further efforts to predict vaccine effectiveness and identify mechanisms of protection.

## Results

### Mass cytometry identifies vaccine-induced CD4 and CD8 ICOS+CD38+CXCR5-subsets

The effect of BNT162b2 immunization was first explored on recipient T cell populations in a cohort of ten healthy donors who had not been previously infected with SARS-CoV-2. Donors had an average age of 41.8 ± 6.3 years. Six donors were male, and nine donors identified as having Caucasian ancestry. By collecting longitudinal peripheral blood samples before immunization, one week following booster immunization (day 28), and at an additional time point approximately three months later (day 105), we captured signatures of the initial immune response to BNT162b2 as well as lasting immunity (**Supplemental Figure 1A**). T cell populations in the pre-vaccination and post-boost samples were initially examined by mass cytometry using a Helios cytometry by time-of-flight mass spectrometry (CyTOF) instrument and a panel of antibodies focused on T cell immune and metabolic phenotypes (**Supplemental Table 1**). Cell density plots of concatenated data of T cells from the pre-vaccination and post-boost samples were first visualized using t-SNE dimensionality reduction (**Figure 1A**). Because MHC-peptide tetramer staining reagents were not available to directly identify expanding antigen specific S protein-reactive CD4 and CD8 T cells, data were analyzed using the recently developed T-REX machine learning algorithm (*24*). By comparing changes in small k-nearest neighbor cell groupings, T-REX specifically identifies populations of cells with the greatest degrees of expansion or contraction from pre- to post-vaccination. In the case of viral infections, these expanding populations were preferentially enriched for virus-specific cells (*24*). This approach sets aside the majority of peripheral blood T cells, which were unchanged, to instead focus on those populations of T cells in phenotypically distinct regions whose abundance increased or decreased by ≥95% in the initial 7 days following vaccination.

**Figure 1.**
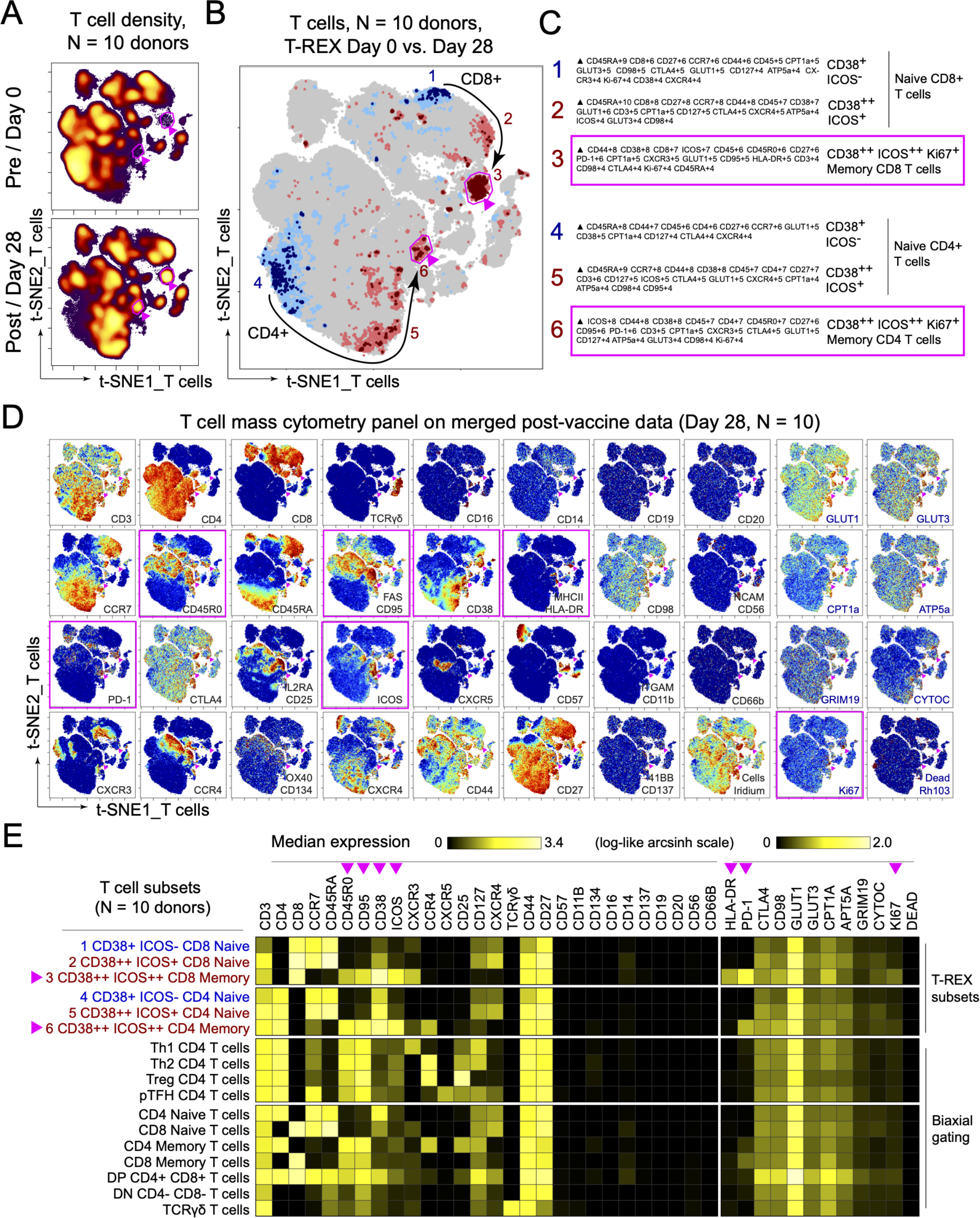
Immune phenotyping of BNT162b2 responding T cells. PBMCs collected from study participants pre-vaccination (day 0), day 28 post-vaccination (7 days post-boost) were analyzed by mass cytometry and data were concatenated for analysis. **A.** CD3+ T cells from 10 donors were pooled into two sets, one for pre-vaccination (taken at day 0) and one for post-vaccination (day 28). Cell density for each set is shown on the t-SNE axes. **B.** T-REX analysis of the CD3+ T cells from the 10 donors is shown. A central t-SNE, performed only using cell surface markers, shaded by T-REX change depicts phenotypically similar cells in regions of great expansion (dark red, ≥95% from post-vaccination day 28) or great contraction (dark blue, ≥95% from pre-) over time following SARS-CoV-2 vaccination. Red or blue population interpretations are shown for major expanding or contracting populations identified by T-REX. **C.** MEM labels show enriched protein features for several populations of great expansion or contraction. All measured features were included in MEM enrichment labels, which only show features enriched by at least +4 on a scale from 0 to 10. Pink boxes are around the MEM labels for the CD4+ and CD8+ memory T cell clusters that are greatly enriched for ICOS and CD38 protein and expanded greatly following vaccination. **D.** Each protein marker is shown on t-SNE axes, with proteins that were enriched on CD4+ and CD8+ ICOS+CD38+ cells in pink boxes. A rainbow intensity scale indicates expression levels with red representing high and dark blue representing low expression. Protein names in blue indicate functional features that were not used in t-SNE analysis, including metabolic markers, Rhodium, and Ki67. Surface proteins in black were used in t-SNE analysis. **E**. Heat maps show all markers measured by mass cytometry for cell populations as determined by T-REX or expert gating. Cell labels in red were defined by T-REX as expanding and blue were defined as contracting by T-REX. Black cell labels were expert gated. Protein markers enriched in CD4+ and CD8+ ICOS+CD38+ cells are indicated with pink arrows.

T-REX revealed related populations of CD4 and CD8 T cells that expanded by ≥95% following vaccination and one population each that contracted by ≥95% (**Figure 1B**). Across the donor cohort, changes in abundance of these cell populations were widely, but not universally, shared in T-REX of individual donor samples (**Supplemental Figure 2A**). Marker Enrichment Modeling (MEM) and specific antibody staining patterns established the protein marker expression patterns characteristic of each population (**Figure 1C**, **Figure 1D**). The most expanded populations were CCR7-CD45R0+ CD4 and CD8 T cell populations that were negative for CXCR5, positive for PD-1, and highly co-expressed CD38 and ICOS (**Figure 1E**). Consistent with extensive expansion, these cells were the most proliferative cell subset based on Ki67 positivity, and were highly metabolically active, based on co-expression of transporters for glucose (GLUT1), amino acids (CD98), and lipids (CPT1a). These cells, thus, reflect non-canonical activated memory T cells. The T cell populations with decreasing abundance, in contrast, were CD45RA+ ICOS- and phenotypically characterized as naïve.

### ICOS+CD38+CXCR5-non-canonical T cell subsets respond specifically to SARS-CoV-2 S protein antigen

The selective enrichment of PD1+ICOS+CD38+CXCR5-CD4 and CD8 T cells following vaccination suggested these cells may specifically recognize viral S protein. To test SARS-CoV-2 specificity and further characterize the CD38+ICOS+ memory T cell populations, fluorescence flow cytometry and FACS were used to characterize cell subsets. Similar to mass cytometry, ICOS+CD38+ CD4 and CD8 cells were present in pre-immunized samples as approximately 1-2% of each T cell subset (**Figure 2A**). Importantly, these cell populations expanded in the post-boost sample and returned to initial frequencies at a later sample collection point. The expanded post-boost populations were also enriched for CCR7-cells (**Figure 2B**) consistent with mass cytometry findings, but not in ICOS^lo^CD38^lo^ T cells from the same samples. To investigate the functional capabilities of ICOS+CD38+ T cells, samples were stimulated with PMA and ionomycin and cytokines were measured. At all collected timepoints, ICOS+CD38+ T cells produced significantly greater levels of IFN-γ and TNFα than ICOS-CD38-cells based on intracellular cytokine flow cytometric staining (**Figure 2C**).

**Figure 2.**
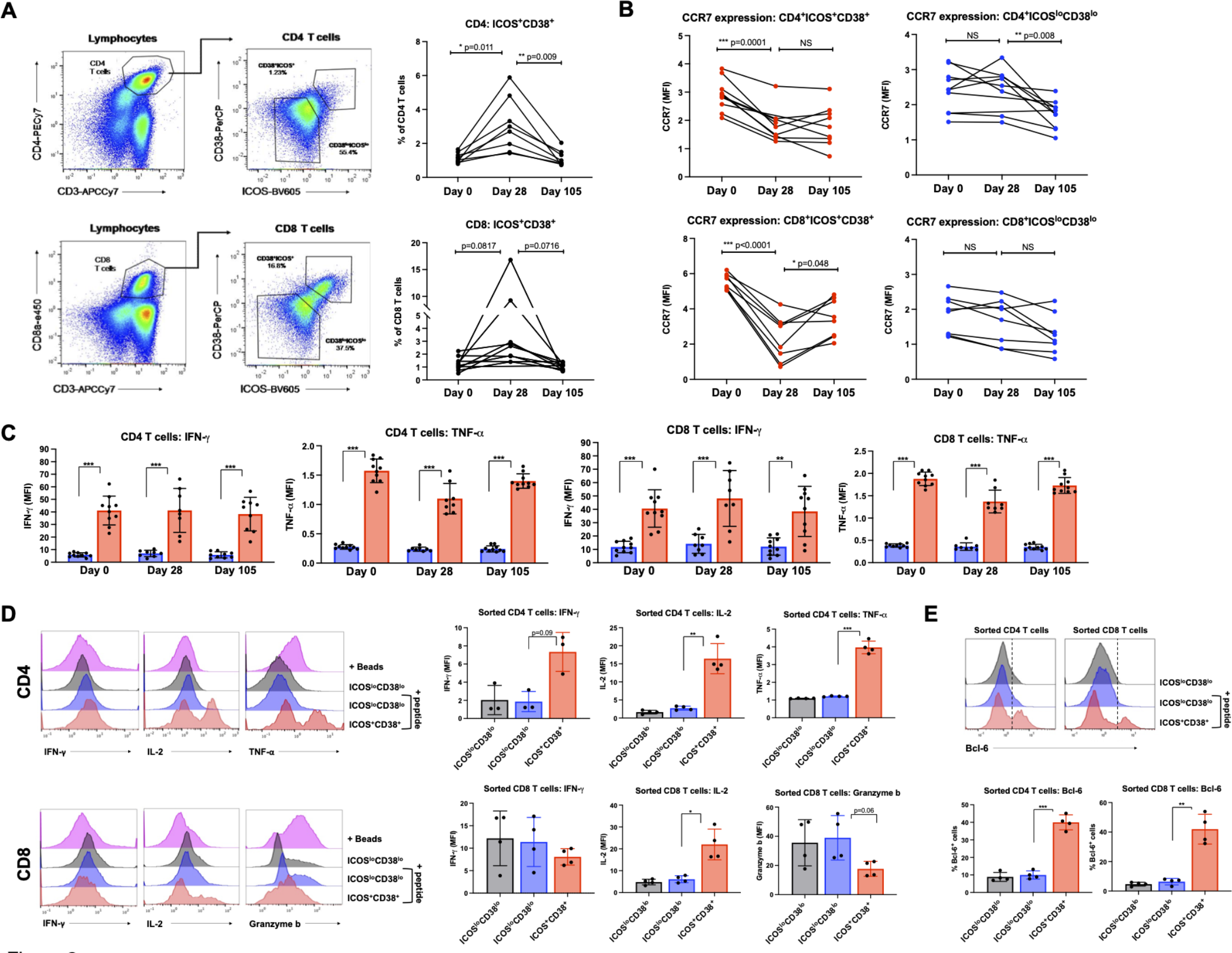
Functional analysis of CD38+ICOS+ CD4 and CD8 T cells and response to SARS-CoV-2 Spike antigen. PBMCs collected from study participants pre-vaccination (day 0), day 28 post-vaccination (7 days post-boost), and day 105 post-vaccination were examined for CD38, ICOS, and CCR7 expression by fluorescence flow cytometry. **A.** CD4 T cells were gated from lymphocytes and defined as ICOS+CD38+ or ICOSloCD38lo. Percentages of ICOS+CD38+ CD4 T cells were quantified and data from each participant is connected by lines. **B.** Mean fluorescence intensity (MFI) of CCR7 within ICOS+CD38+ CD4 T cells or ICOSloCD38lo CD4 T cells. CD8 T cells were gated from lymphocytes and defined as ICOS+CD38+ or ICOSloCD38lo. Percentages of ICOS+CD38+ CD8 T cells were quantified from each donor. Mean fluorescence intensity (MFI) of CCR7 within ICOS+CD38+ CD8 T cells or ICOSloCD38lo CD8 T cells. Significance was determined by paired ANOVA mixed-effects model. NS= not significant. **C.** PBMCs were stimulated with PMA/ionomycin and examined for cytokine production. T cells defined as ICOS+CD38+ (red) or ICOSloCD38lo (blue) were examined for IFN-γ (left) or TNFα (right). **D.** Day 28 PBMC samples were sorted based on CD38 and ICOS expression. T cells were labeled with CellTrace Violet (CTV) and stimulated with autologous (day 0 CD3-depleted) PBMCs with or without SARS-CoV-2 Spike peptide pool (+ peptide, blue and red). CTV-labeled Cells were analyzed after 48h of stimulation for cytokine production and effector function. Sorted CD4 T cell populations were analyzed for IFN-γ, IL-2, and TNFα. Sorted CD8 T cell populations were analyzed for IFN-γ, IL-2, and Granzyme b. **E.** BCL6 expression in sorted T cell populations, stimulated with peptide. Significance determined by paired ANOVA. * p<0.5, ** p<0.005, ***p<0.0005

To evaluate the antigen specific T cell response to SARS-CoV-2 S antigen, PBMCs from each time point were incubated with recombinant SARS-CoV-2 S protein or T cell activation beads and cytokine responses were measured in culture supernatant. Interestingly, post-boost samples clustered into responders and non-responders for IFN-γ and TNFα production (**Supplemental Figure 3A, B**). IL-2, IL-4 and IL-17a were unaffected by stimulation of T cells with S protein (**Supplemental Figure 3C-E**). Furthermore, a subset of samples from all three time points produced IL-6 in response to S protein, suggesting cross-reactivity with prior coronavirus exposures (**Supplemental Figure 3F**). Especially notable were the late collection time points in which three donor samples produced more IL-6 when incubated with S protein than positive control polyclonal CD3/2/28 bead stimulated samples. Samples from day 105 also showed an increase in the frequency of IL-2 and IFN-γ producing CD8 T cells when restimulated with S protein after 11 days of culture (**Supplemental Figure 3G**).

To directly test antigen specificity and further characterize the expanding T cells identified by T-REX, ICOS+CD38+ and ICOS^lo^CD38^lo^ cells were isolated from four donor samples using FACS and protein markers from T-REX populations. Cells were then labeled with CellTrace Violet and stimulated with CD3-depleted autologous PBMCs with or without SARS-CoV-2 S peptide pool (**Supplemental Figure 3H**). ICOS+CD38+ CD4 T cells produced IFN-γ, IL-2, and TNFα in response to SARS-CoV-2 S peptide stimulation, while peptide-stimulated ICOS^lo^CD38^lo^ T cells from the same sample failed to produce cytokines (**Figure 2D**). CD8 ICOS+CD38+ T cells also produced significant IL-2 in response to peptide stimulation, but had decreased Granzyme B, suggesting potential post-activation degranulation. Both CD4 and CD8 ICOS+CD38+ populations showed evidence of metabolic reprogramming, as mTORC1 pathway activity as shown by levels of phospho-S6 trended higher in each donor (**Supplemental Figure 3I**). Expression of the glucose transporter GLUT1, however, was unchanged or modestly reduced (**Supplemental Figure 3J**). Although the ICOS+CD38+ CD4 and CD8 T cells lacked CXCR5, the expanding S protein-specific cells shared some characteristics with circulating T follicular helper cells (cTfh) as a portion of these cells expressed the Tfh characteristic transcription factor BCL6 (**Figure 2E**).

### BNT162b2 induces expansion of CD38+CD43+ plasmablasts

The B cell response to BNT162b2 was next examined using a similar approach as applied to the T cell response. Peripheral blood samples from the same donor cohort were analyzed by mass cytometry using a panel of antibodies focused on B cell populations and metabolism. T-REX analysis of equally sampled B cells from all individuals showed that while most B cell populations did not change following boost, there were 9 phenotypically-distinct areas of increased or decreased cell abundance (**Figure 3**). Of these, seven were populations that expanded and two were populations that contracted. The populations that expanded ≥95% from day 0 to day 28 included activated IgM+ B cells, memory B cells, and plasmablasts, while naïve B cells greatly contracted over this time. While individual donors showed some heterogeneity, these populations remained broadly evident (**Supplemental Figure 4**). Using MEM to characterize cells identified by T-REX, the IgM-IgD-plasmablast population was further defined as CD20-CD38+CD43+ and metabolically active, with high expression of the iron transporter CD71, CD98, and Cytochrome C (**Figure 3C-E**).

**Figure 3.**
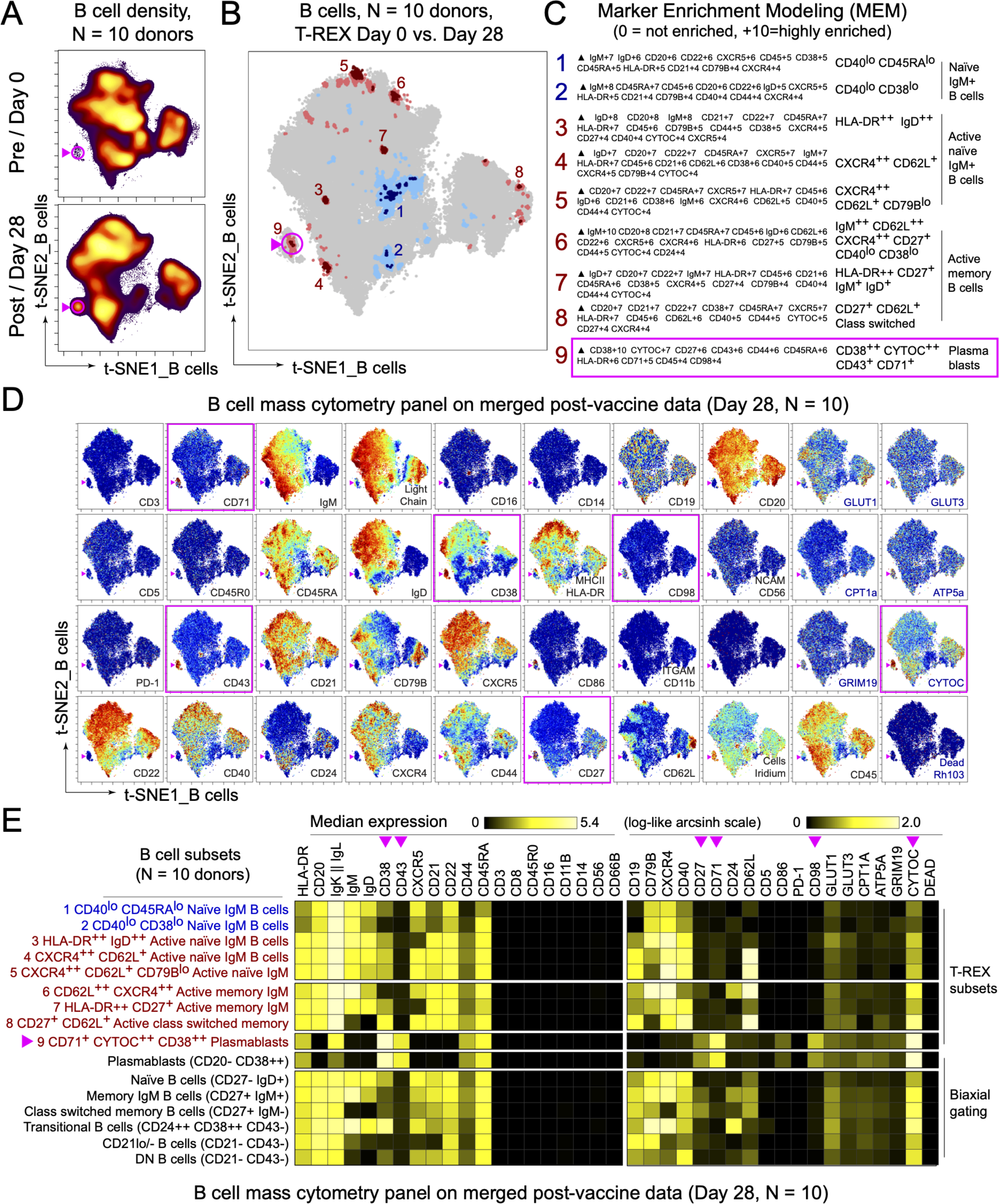
Immune phenotyping of BNT162b2 responding B cells. PBMCs collected from study participants pre-vaccination (day 0), day 28 post-vaccination (7 days post-boost) were analyzed by mass cytometry and data were concatenated for analysis of expanded B cell subsets and plasmablasts. **A.** B cells from 10 donors were pooled into two sets, one for pre-vaccination (taken at day 0) and one for post-vaccination (day 28). Cell density for each set is shown on the t-SNE axes. **B.** T-REX analysis of the B cells from the 10 donors is shown. A central t-SNE, performed only using cell surface markers, shaded by T-REX change depicts phenotypically similar cells in regions of great expansion (dark red, ≥95% from post-vaccination day 28) or great contraction (dark blue, ≥95% from pre-) over time following SARS-CoV-2 vaccination. Red or blue population interpretations are shown for major expanding or contracting populations identified by T-REX. **C.** MEM labels show enriched protein features for several populations of great expansion or contraction. All measured features were included in MEM enrichment labels, which only show features enriched by at least +4 on a scale from 0 to 10. A pink box is around the MEM label for the plasmablast cluster that expanded greatly following vaccination. **D.** Each protein marker is shown on t-SNE axes, with proteins that were enriched on the plasmablast population in pink boxes. A rainbow intensity scale indicates expression levels with red representing high and dark blue representing low expression. Protein names in blue indicate functional features that were not used in t-SNE analysis, including metabolic markers, Rhodium, and Ki67. Surface proteins in black were used in t-SNE analysis. **E.** Heat maps show all markers measured by mass cytometry for cell populations as determined by T-REX or expert gating. Cell labels in red were defined by T-REX as expanding and blue were defined as contracting by T-REX. Black cell labels were expert gated. Protein markers enriched in plasmablasts are indicated with pink arrows.

### Serologic response to BNT162b2 does not correlate to pre-existing immunity for endemic coronaviruses

BNT162b2-induced serological responses were measured by testing donor plasma samples from day 0, day 28, and day 105 for reactivity to S proteins from SARS-CoV-2 Wuhan-1, SARS-CoV-2 Beta, SARS-CoV-2 Alpha, HCoV-OC43, and HCoV-HKU1. Robust IgG responses were measured against SARS-CoV-2 Wuhan-1 by ELISA, with more varied IgM responses (**Figure 4A**, **Supplemental Figure 5A**). These patterns were generally consistent across the S proteins from circulating variants of concern (VOCs). Interestingly, an IgA vaccine response was also evident in most donors. Most donors also displayed signatures of pre-existing immunity to the coronaviruses HCoV-OC43 and HCoV-HKU1, consistent with their endemic nature among the human population(*27*). Antibody levels to these endemic strains, however, did not change following BNT162b2 vaccination. Next, using a replication-competent vesicular stomatitis virus (VSV) SARS-CoV-2 (*28*), Real Time Cell Analysis assay(*29*) was performed to evaluate the neutralization activity of donor plasma at both pre- and at multiple timepoints post-vaccination. While pre-vaccine samples exhibited no SARS-CoV-2 neutralization activity, most donors showed some neutralizing activity 3 months post-boost, although 3 of the 10 donors showed only weak neutralization at that timepoint (**Figure 4B**). Interestingly, a sharp decline in neutralization potency was observed for samples 3 months post-boost compared to day 7-8 post-boost in 3 of the 4 donors with multiple timepoints post-vaccination. Despite a modest trend for HCoV-HKU1 IgG, pre-existing antibody titers to HCoV-OC43 and HCoV-HKU1 did not correlate with SARS-CoV-2 neutralization potency 3 months post-vaccination (**Figure 4C**). Among the serological variables that were tested, SARS-CoV-2 IgG level 3 months post-boost was the only one with a significant correlation with VSV SARS-CoV-2 neutralization potency (**Figures 4C and 4D**). Therefore, pre-existing coronavirus antibody did not appear to be a major determinant for defining the antibody response to BNT162b2 vaccination.

**Figure 4.**
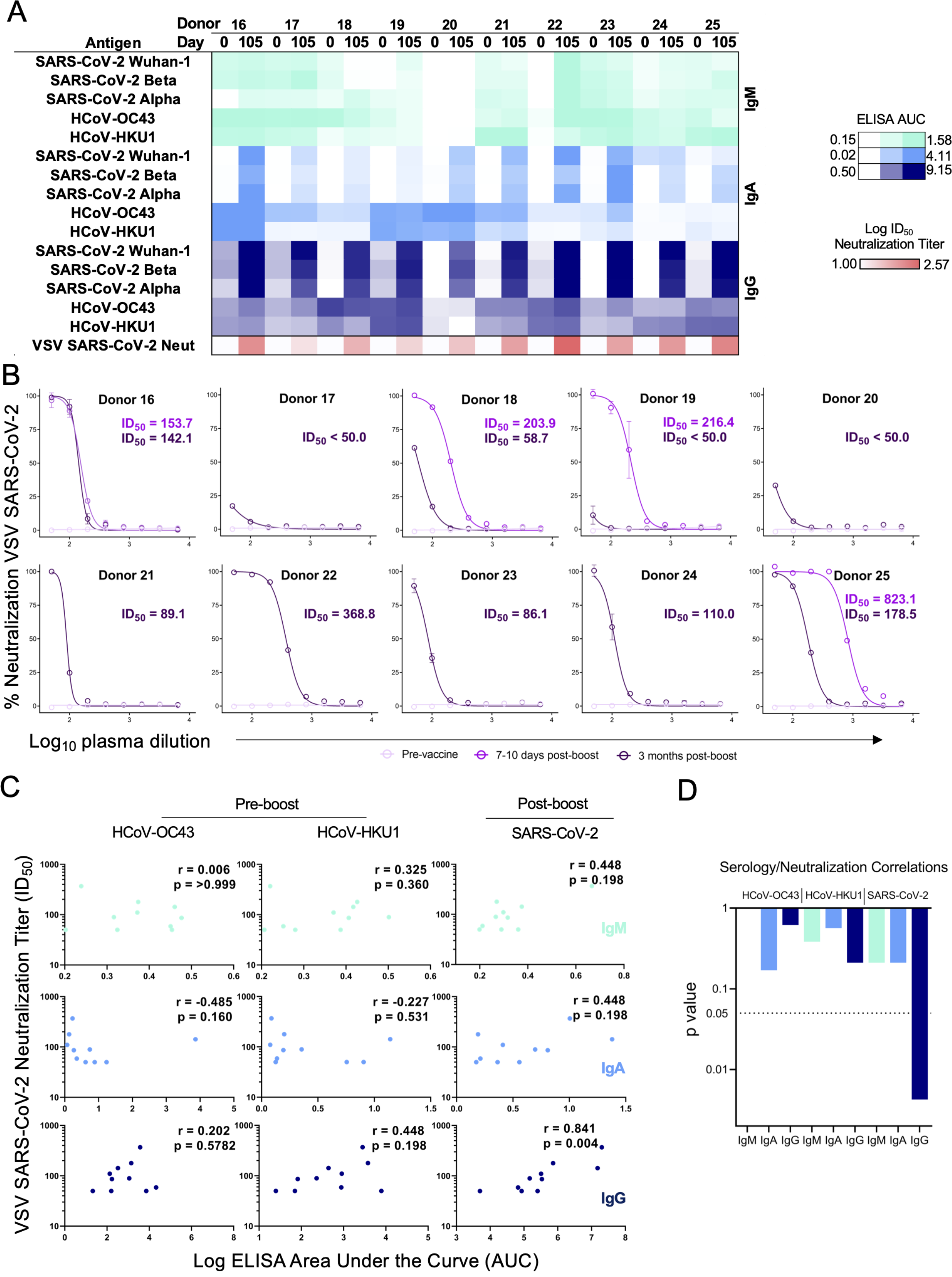
Serologic response induced by BNT162b2. Sera were analyzed from healthy donors pre-vaccination and at a late time point. **A.** ELISA area under the curve (AUC) values from vaccine recipient plasma are depicted as a heatmap against spike proteins from SARS-CoV-2 Wuhan-1, SARS-CoV-2 Alpha, SARS-CoV-2 Beta, HCoV-OC43, and HCoV-HKU1 coronaviruses. The minimum signal is represented in white and maximum signal is depicted by teal, blue, and navy blue for IgM, IgA, and IgG, respectively. At the bottom of the serology heatmap, Log ID_50_ neutralization titer is depicted from zero neutralization (white) to the maximum Log ID_50_ value quantified in the donor cohort. **B.** VSV SARS-CoV-2 neutralization curves of vaccine recipient plasma. Pre-vaccine, 7-10 days post-boost (collected only from donors 16, 18, 19, and 25), and 3 months post-boost are depicted in light pink, pink, and purple, respectively. %Neutralization (y-axis) is plotted as a function of plasma dilution (x-axis). **C.** Spearman correlations for serological responses as log ELISA AUC (x-axis) of patient cohort against pre-vaccination HcoV-OC43 (left), HcoV-HKU1 (middle), and post-boost SARS-CoV-2 (right) as a function of neutralization titer expressed as ID_50_ (y-axis). IgM is depicted in teal (top), IgA is shown in blue (middle), and IgG is shown in navy blue (bottom). The respective rho and p values are shown in the top right of each plot. **D.** Individual statistical significance comparison values are depicted as a bar graph for HCoV-OC43, HCoV-HKU1, and SARS-CoV-2 (left to right). IgM is depicted in teal, IgA is shown in blue, and IgG is shown in navy blue.

### Antigen selectivity of the SARS-CoV-2-reactive B cell response to BNT162b2 increases over time

While numerous antigen-specific antibodies have been isolated from SARS-CoV-2 infection (*30-32*) and more recently from vaccination (*33*), little is currently known about the antigen specificities of individual B cells within the repertoire of SARS-CoV-2 vaccinees and how these change over time. We thus set out to characterize the evolution of the SARS-CoV-2 specific B cell repertoire by analyzing multiple time points pre- and post-vaccination of a single healthy male Caucasian donor of age 45-50 with no prior history of SARS-CoV-2 infection (**Supplemental Figure 1B**) with LIBRA-seq (*25, 34*). LIBRA-seq enables high-throughput mapping of B cell receptor sequence to antigen specificity for large numbers of B cells per sample against a diverse set of coronavirus antigens. The antigen specificity of the polyclonal plasma from this donor was largely similar to what was observed for the ten-donor cohort described above, with BNT162b2 immunization leading to the emergence of SARS-CoV-2-reactive IgM, IgA, and IgG antibodies (**Figure 5A**). While some reactivity with the closely related coronavirus SARS-CoV was observed, little reactivity was observed against the more distantly related MERS-CoV. Although SARS-CoV-2 S-reactive IgA and IgG were present at day 14 post-vaccine prime, neutralizing antibody titers were first detected at day 28 (7 days post-boost) at levels that remained largely unchanged at day 42 (**Figure 5B**).

**Figure 5.**
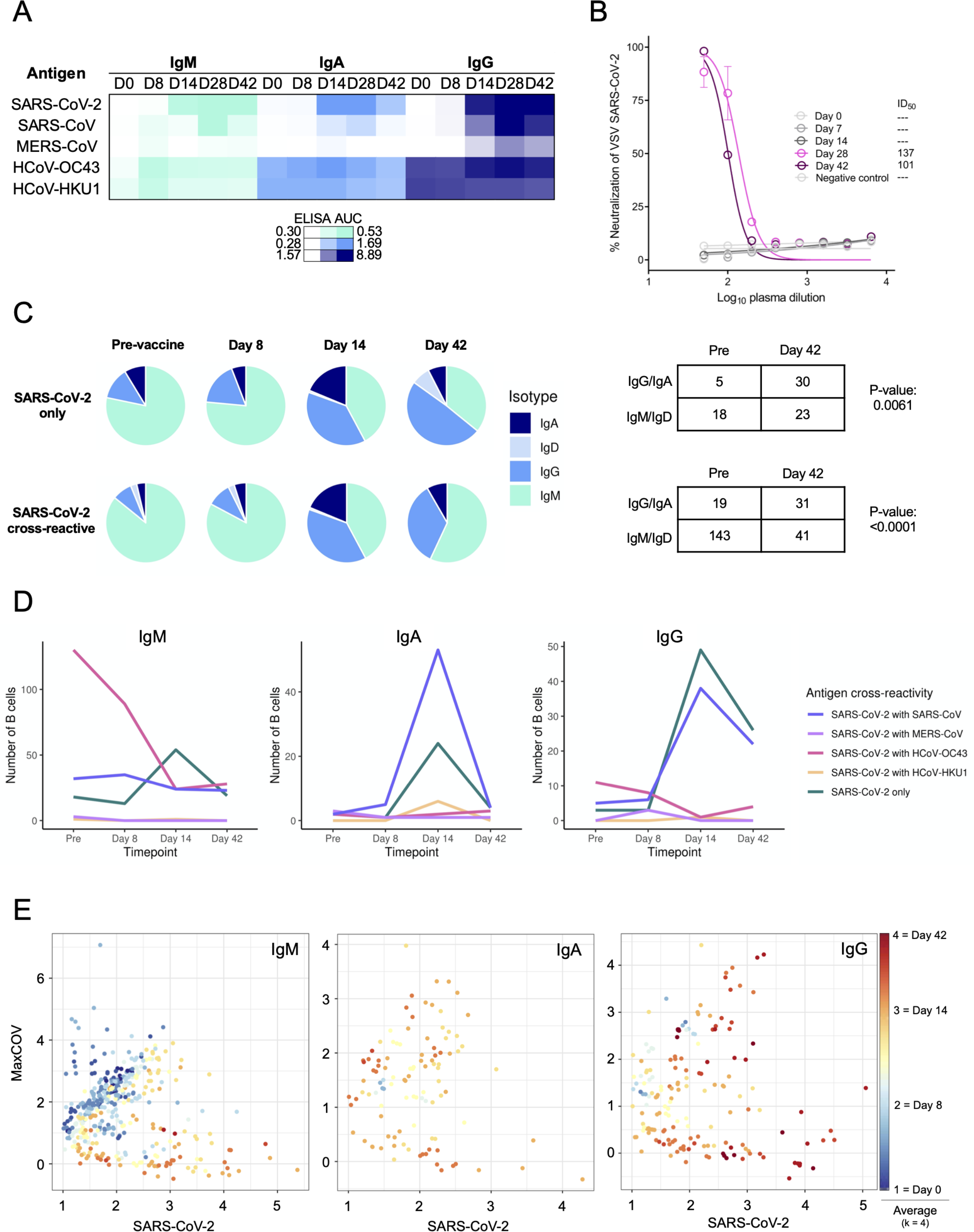
LIBRA-seq characterization of the antigen specificity of the SARS-CoV-2-reactive B cell response to BNT162b2. A single donor with multiple longitudinal samples was analyzed serologically and by LIBRA-seq. **A.** Plasma ELISA area under the curve (AUC) values are depicted as a heatmap against spike proteins for SARS-CoV-2, SARS-CoV, MERS-CoV, HCoV-OC43, and HCoV-HKU1. The minimum signal is represented in white and maximum signal is depicted by teal, blue, and navy blue for IgM, IgA, and IgG, respectively. **B.** VSV SARS-CoV-2 neutralization of longitudinal timepoints. % Neutralization (y-axis) is plotted as a function of plasma dilution (x-axis). The negative control sample, Day 0, Day 8, and Day 14 timepoints are depicted in grey, with Day 28 and Day 42 curves shown in pink and purple respectively. The inhibitory dose at 50% neutralization (ID_50_) values of each timepoint are denoted to the right of graph. **C.** Pie charts representing: (top) SARS-CoV-2 specific B cells with an associated LIBRA-seq score ≥1 for SARS-CoV-2; and (bottom) SARS-CoV-2 cross-reactive B cells (bottom) with an associated LIBRA-seq score ≥1 for SARS-CoV-2 and for at least one other coronavirus antigen (MERS-CoV, SARS-CoV, HCoV-OC43 or HCoV-HKU1). For the pre-vaccine, day 8, day 14, and day 42 timepoints, the segments in each pie chart represent the number of antibody sequences with the isotypes IgD (light blue), IgM (teal), IgG (blue), IgA (navy blue). Also shown is a statistical comparison of isotype distribution of SARS- CoV-2 specific (top) and SARS-CoV-2 cross-reactive (bottom) B cells in the pre-vaccine and day 42 post-vaccination timepoints. The values in each table represent the number of antibody sequences with the designated isotypes. **D.** Evolution of cross-reactive SARS-CoV-2 and SARS-CoV-2-only B cells from each isotype (separate plots) over time. Each line shows the number of B cells (y-axis) for either SARS-CoV-2-only (blue) or SARS-CoV-2 cross-reactive with other coronaviruses (designated colors) B cells at the four timepoints (x-axis). **E.** Individual IgM, IgA, and IgG expressing cells graphed for cumulative cross-reactive (Max COV) and SARS-CoV-2 LIBRA-seq scores shaded based on k=4 nearest neighbor averaging for time of sample collection.

Using a LIBRA-seq antigen library consisting of DNA oligo-tagged S proteins from SARS-CoV-2, SARS-CoV-2 D614G, SARS-CoV, MERS-CoV, HCoV-HKU1, HCoV-OC43, as well as the negative control proteins, B cells were enriched for antigen-positive cells by FACS (**Supplemental Figure 6A**) and mapped to their antigen reactivity profile utilizing a next-generation sequencing readout. Clonal lineage tracing of BCR sequences from each timepoint identified several clusters containing sequences with high LIBRA-seq scores for SARS-CoV-2 (>1) across multiple timepoints (**Supplemental Figure 6B**). LIBRA-seq identified both B cells that were specific to SARS-CoV-2 (henceforth referred to as SARS-CoV-2-only B cells) as well as B cells that were cross-reactive between SARS-CoV-2 and other coronaviruses (cross-reactive B cells) **(Supplemental Figure 7)**. Both SARS-CoV-2-only and cross-reactive B cells showed a shift from IgM to IgG after vaccination, suggesting antigen-driven class switching (**Figure 5C**). IgM remained more prevalent than IgG in the cross-reactive B cell population, suggesting a stronger class-switching potential for SARS-CoV-2-only over cross-reactive B cells in response to vaccination. SARS-CoV-2 cross-reactivity was primarily observed in HCoV-OC43-reactive IgM cells in this donor, although the overall levels of SARS-CoV-2 cross-reactive B cells generally decreased at later timepoints post-vaccination (**Figure 5D**). In contrast, the levels of SARS-CoV-2-only IgG+ B cells and IgG+ B cells that cross-reacted between SARS-CoV-2 and the closely related SARS-CoV were substantially increased at days 14 and 42. Interestingly, while the levels of SARS-CoV-2-reactive B cells peaked at day 14 for both the IgG and IgA isotypes, the IgA levels were reduced to near baseline at day 42, while IgG levels at day 42 were decreased but clearly present. Changes in B cell cross-reactivity and isotype abundance were also evident when analyzed at the single-cell level using antigen-specific LIBRA-seq scores as a metric for comparing antigen specificity evolution (**Figure 5E**). These results suggest an evolution in SARS-CoV-2 antibody specificity and cross-reactivity in response to BNT162b2 vaccination, supporting a model of immune progression from early prevalence of endemic cross-reactive IgM to more highly selective SARS-CoV-2 IgG production over time.

### SARS-CoV-2-binding memory B cells and plasmablasts expand following BNT162b2 vaccination

The LIBRA-seq B cell receptor sequence and antigen specificity data was further combined with RNA-seq to identify gene expression patterns and B cell subset identities of SARS-CoV-2-binding B cells over time following vaccination. Seurat and LIBRA-seq analyses were integrated to enable immune repertoire analysis within transcriptionally-defined B cell subsets. Nine clusters were identified by UMAP across all B cells based on gene expression profiles that represented naive, predominantly IgM memory (termed “memory”), predominantly IgA or IgG-switched memory (termed “switched memory”), or plasmablast populations (**Figure 6A, B**). Characteristic gene expression patterns defined each B cell subset (**Figure 6C**, **Supplemental Figure 8A**). Only a modest number of genes were differentially expressed in LIBRA-seq predicted SARS-CoV-2-binding B cells relative to other members of those cell clusters (**Supplemental Figure 8B**). These did include, however, selectively elevated *CCR9*, *IgHM*, and *IGLC1* and decreased Galectin 1 (*LGALS1*) in the SARS-CoV-2-binding plasmablast populations, suggesting active migration and antibody production. When combined with LIBRA-seq scores for putative antigen specificity, predicted SARS-CoV-2-binding B cells expressed a wide range of *IGHV* gene segments (**Supplemental Figure 8C**). Class-switching is associated with protective immune responses, thus further analyses focused on class-switched SARS-CoV-2-binding B cells. Consistent with plasmablast populations identified by T-REX analysis of B cell protein expression (**Figure 3**), LIBRA-seq suggested SARS-CoV-2-binding B cells were represented among both switched memory and plasmablast cell clusters (**Figure 6D**). Analysis of class-switched, SARS-CoV-2-binding B cells revealed the majority had no or only modest degrees of somatic hypermutation, which did not increase with time (**Figure 6E**, **Supplemental Figure 8D**). By day 14 a robust SARS-CoV-2 binding plasmablast population with limited somatic hypermutation was evident that diminished by day 42 after initial BNT162b2 vaccination (**Figure 6F**). These plasmablasts did not have significant LIBRA-seq cross-reactivity scores for endemic coronaviruses but included plasmablasts selective for SARS-CoV-2 or with dual selectivity for SARS-CoV-2 and SARS-CoV (**Figure 6G**).

**Figure 6.**
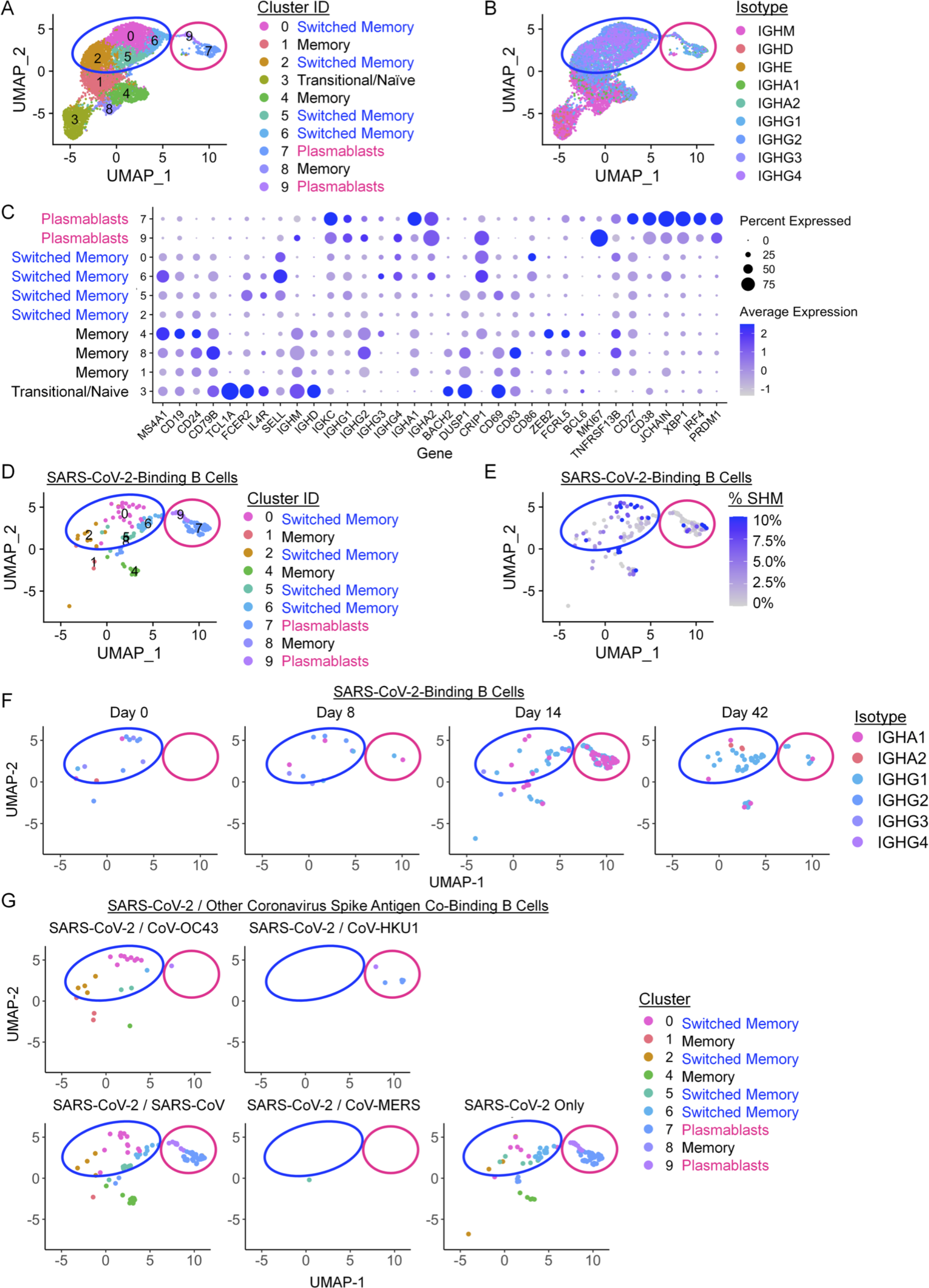
BNT162b2 vaccination drives IgG and IgA-switched SARS-CoV-2-binding memory B cell and plasmablast expansion. B cells that bound a panel of LIBRA-seq antigens were purified from peripheral blood samples and single cell RNA-seq/BCR-seq/LIBRA-seq data were integrated. Cells were scored positive for binding a given antigen for LIBRA-seq scores > 1. **A.** UMAP identified clusters of B cells based on transcriptional similarity. Antibody-secreting cells (pink circle encompassing clusters 7 and 9) and isotype-switched memory B cells (blue circle encompassing clusters 0, 2, 5, and 6) are highlighted across all UMAP plots. **B.** BCR isotypes are shown among sorted B cells. **C.** Selected gene expression profiles within each cluster are shown. **D-F.** SARS-CoV-2-binding B cells were identified using LIBRA-seq and non-class-switched B cells were filtered out for detailed analysis. **D.** UMAP identified clusters of SARS-CoV-2-binding B cells based on transcriptional similarity. **E.** IgH somatic hypermutation frequency of SARS-CoV-2-binding B cells is indicated. **F.** SARS-CoV-2-binding B cell isotypes are shown for each time point. **G.** SARS-CoV-2-binding B cells are shown, each panel indicates cells which also bound the indicated coronavirus S antigens.

### Integrated associations of antigen-specific cell and antibody responses to BNT162b2

To test associations of the antigen-specific CD4, CD8 T and B cell plasmablast populations and the antibody response to BNT162b2, the abundance of ICOS+CD38+ CD4 or CD8 and plasmablast cell populations were compared across donors (**Figure 7A**). The CD4, CD8, and plasmablast populations were associated with each other to support a coordinated response. Further, plasmablast frequencies showed a trend towards significance and ICOS+CD38+ CD8 T cells correlated with eventual levels of SARS-CoV-2 specific IgG. The three donors with the lowest levels of neutralizing antibodies (red and pink) also had the lowest numbers of ICOS+CD38+ CD8 cells. Intriguingly, donor 17 (red) had the lowest abundance of ICOS+CD38+ CD8 cells and plasmablasts and was subsequently infected with SARS-CoV-2. While donor 17 represents a single case, the observation supports the idea that BNT162b2-induced expansion of non-canonical memory ICOS+CD38+ CD4 and CD8 T cells may be critical to drive plasmablast expansion, antibody production, and protective immunity to SARS-CoV-2.

**Figure 7.**
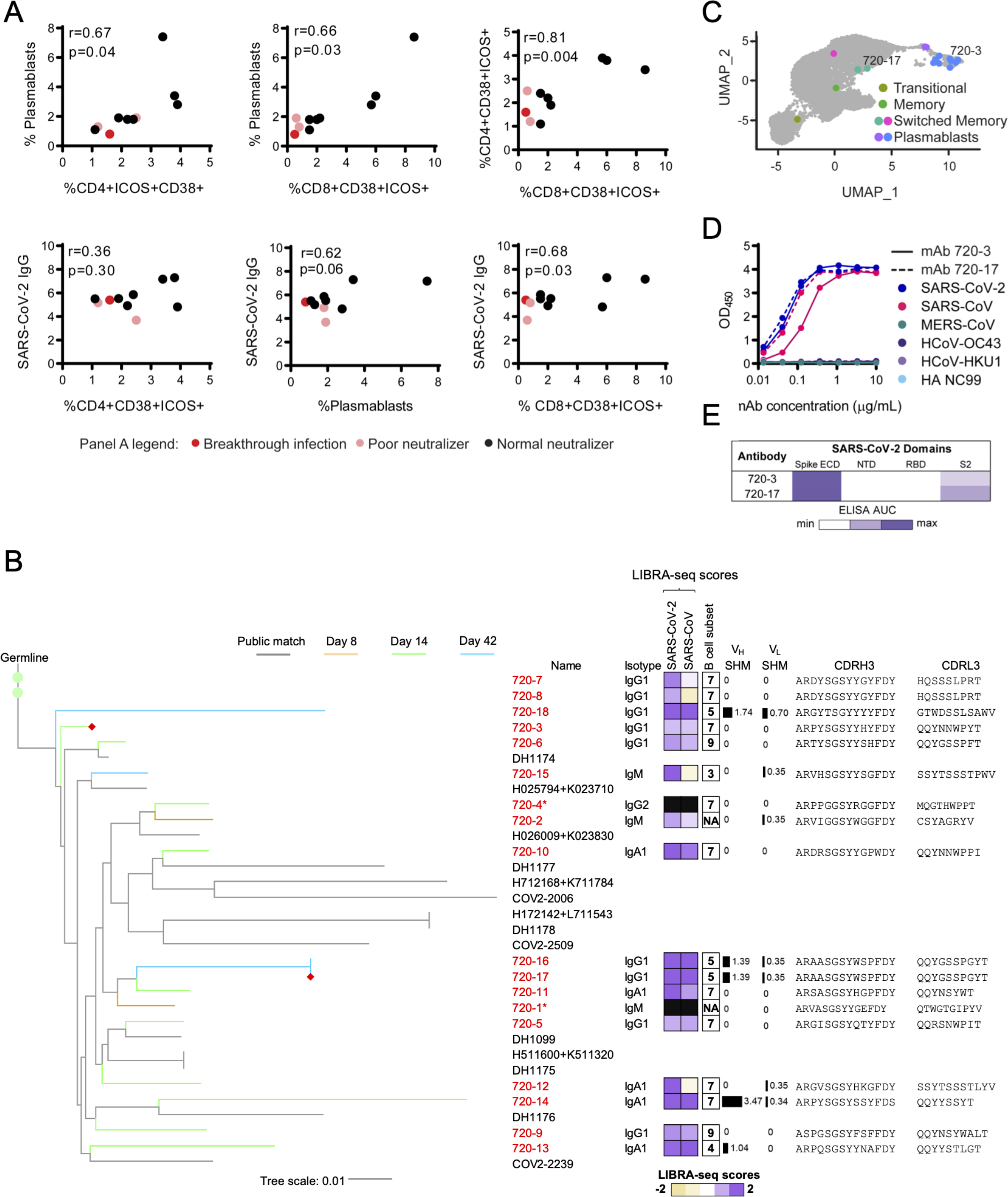
Associations of antigen-specific cell populations and antibody responses. A. Correlation of biaxially gated plasmablast and T cell populations with other cellular subsets and SARS-CoV-2 IgG using Pearson correlations. Patients with normal neutralization titers are shown in black, poor neutralizers with no known breakthrough infection are shown in pink, and a poor neutralizer with known breakthrough infection is shown in red. **B.** Phylogenetic tree of a public antibody cluster comprised of LIBRA-seq-identified sequences (antibody names: red; branches: colored by timepoint) and previously published SARS-CoV-2 antibody sequences from the CoV-AbDab database (grey), with recombinantly expressed antibodies 720-3 and 720-17 highlighted with a red diamond at the end of their respective branches. For each sequence from the cluster, sequence features including isotype, LIBRA-seq scores, % SHM of V-gene, and amino acid sequences of CDRH3 and CDRL3 are displayed. Nucleotide level percentage of SHM changes of V-genes of both heavy and light chains were reported as bars with numerical values. LIBRA-seq scores are shown as a range with tan-white-purple representing -2 to 0 to 2. Scores higher or lower than this range are shown as -2 and 2, respectively. The B cell subsets are classified by scRNA-seq analysis. **C.** Location of B cells in cluster 720 including recombinantly expressed monoclonal antibodies (720-3 and 720-17) on UMAP identified clusters of B cells based on transcriptional similarity. **D.** ELISAs for antibodies selected for monoclonal antibody characterization are shown against spike proteins for SARS-CoV-2 (black), SARS-CoV (pink), MERS-CoV (teal), HCoV-OC43 (purple), HCoV-HKU1 (light purple), and HA NC99 (light blue) for antibodies 720-3 (solid lines) and 720-17 (dashed lines). **E.** Epitope mapping ELISA AUC for antibodies 720-3 and 72017 against SARS-CoV-2 spike ECD, and truncated subunit domains NTD, RBD, and S2.

We next examined the antibody sequence relationships of B cells elicited by vaccination from the longitudinally collected donor. Interestingly, a cluster of antibody heavy chain sequences, referred to as cluster 720, was found to exhibit high similarity with public antibody sequences from the CoV AbDab database (*35*) of individuals with natural COVID-19 infection (**Figure 7B**). Consistent with previous studies, this is indicative of a convergent B cell response from natural infection and vaccination (*36*). Interestingly, however, despite the high similarity in heavy chain sequences, the members of cluster 720 utilized a diversity of light chain V-genes and CDRL3 sequences. The B cells from cluster 720 included sequences with IgM, IgA, and IgG isotypes, and originated from switched memory and plasmablast cell clusters (**Figure 7C**) that displayed only low to moderate degrees of somatic hypermutation (**Figure 7B**). Day 8 sequences from cluster 720 were an IgM isotype, whereas the sequences from days 14 and 42 appeared as IgG and IgA isotypes (**Figure 7B**). To characterize individual antibodies from cluster 720, we expressed heavy- and light-chain pairs as recombinant IgG for two members from diverse branches in the tree, 720-3 (using IGKV3-15) and 720-17 (using IGKV3-20), and tested for reactivity against coronavirus S proteins. Consistent with their LIBRA-seq scores, these antibodies were cross-reactive to SARS-CoV-2 and the closely related SARS-CoV but showed no reactivity to the endemic coronaviruses HCoV-HKU1 and HCoV-OC43 (**Figure 7D**). Epitope mapping experiments revealed that these two antibodies were specific to the S2 domain of spike (**Figure 7E**, **Supplemental Figure 9**).

## Discussion

Here we describe the antigen-specific cellular response and development of the specific antibody repertoire over time in a longitudinal cohort of healthy donor recipients of the BNT162b2 vaccine. Machine learning analyses of proteomic mass cytometry data and single cell sequencing of the SARS-CoV-2 S B cell repertoire identified antigen-specific CD4 T cell, CD8 T cell, B cell and plasmablast populations. PD1+CD38+ICOS+CXCR5-CD4 and CD8 T cells produced TNFα and IFN-γ in response to stimulation with SARS-CoV-2 S protein and shared features of memory or Tfh. The B-cell repertoire shifted from initial apparent cross-reactivity with endemic coronavirus prior to immunization to a more SARS-CoV-2-selective response later, marked by expansion of IgA and IgG plasmablasts. Importantly, antigen-specific cell subsets identified early after boost correlated with sustained IgG antibody response at day 105, and both the CD4 and CD8 populations of ICOS+CD38+ T cells were deficient in a donor who later experienced a breakthrough SARS-CoV-2 infection.

Antigen-specific CD4 and CD8 T cells that drive the vaccine response were phenotypically distinct from canonical peripheral blood T cell populations. SARS-CoV-2 S protein specific CD4+ PD1+CD38+ICOS+CXCR5-did resemble CD38+ICOS+CXCR5-T cells previously identified following SARS-CoV-2 infection (*4*) and a prior study of BNT162b2 vaccination (*37*), but had several previously undescribed characteristics. A subset of these antigen specific cells did express the Tfh transcription factor BCL6, and PD1+ICOS+ cTfh cells have been previously associated with vaccine responses (*38-40*). However, they lacked CXCR5 that is characteristic of Tfh and CD4 T cells produced IL-2 and TNFα when antigen-stimulated. The CD38+ICOS+ antigen specific population resemble rhinovirus-specific tissue homing memory CD4+ T cells that express CCR5 and CD38 and are associated with transcription factor TBET (*41*) and may reflect extrafollicular PD-1+CXCR5-CD4+ T cells observed following viral infections (*42*). Alternatively, these cells may include memory Tfh or IFN-γ-producing Thf1 populations that have down regulated CXCR5 in circulation as they enter a memory pool (*43-45*). CD38+ICOS+CXCR5-cells thus appear as antigen-experienced memory T cell that may be tissue homing and arise from multiple T cell subtypes, such as Th1 and Tfh CD4 T cells to be poised for long-lived anti-viral memory responses.

In addition to induction of SARS-CoV-2 neutralizing antibodies (*46*), cytotoxic T cell responses also contribute to vaccine-mediated protection. CD8+ T cells support life-long immunity against influenza (*47*), EBV(*48*), and CMV (*49*), and induction of a robust CD8+ T cell response is an emerging focus in vaccine development (*50, 51*). Here, we identified antigen-specific CD8+ PD1+CD38+ICOS+CXCR5-cytotoxic memory T cell population capable of cytokine production in response to S antigenic peptides. Similar CD8+ T cells are induced by yellow-fever vaccination (*52*) that can persist up to 25 years (*53*). CD8+ PD1+CD38+ICOS+CXCR5-T cells described here may, therefore, also provide immunological memory for long-lasting protection against SARS-CoV-2. Interestingly, cellular immunity may provide protection to individuals who mounted a suboptimal humoral antibody response and the donor with breakthrough SARS-CoV-2 infection failed to generate CD8+CD38+ICOS+ T cells. If confirmed in a larger cohort, CD8+CD38+ICOS+ T cells may provide a novel marker of successful immunization and protection following SARS-CoV-2 vaccination.

SARS-CoV-2 S protein-specific B cells identified here exhibited similar phenotypic properties as previous studies. These include expression of the B cell activation marker CD71, an iron transport receptor that is commonly expressed on antigen-specific B cell subsets including in the context of vaccination (*53, 54*). Reports of CD71 expression after COVID-19 infection (*55, 56*) are consistent with overlap of mRNA vaccination and natural SARS-CoV-2 infection. Limited pre-existing immunity to endemic coronaviruses prior to SARS-CoV-2 vaccination was detected in this study, albeit at varying levels between donors. Although a previous report noted an increased serological response in recovered COVID-19 individuals (*57*), we noted only modest changes in HCoV-OC43 serology following vaccination. Instead, LIBRA-seq antigen-specificity data showed an initial HCoV-OC43, SARS-CoV-2 cross-reactive IgM-expressing B cell that decreased in frequency over time and evolved to development of IgA- and IgG-expressing B cells with greater SARS-CoV-2 specificity. Given HCoV-OC43 and HCoV-HKU1 utilize different cell receptors for host entry, the lack of correlation between pre-existing coronavirus immunity and SARS-CoV-2 neutralization is not surprising. Instead, antibodies and memory B cells that target the more conserved S2 portion of spike may promote anti-viral function via Fc effector function mechanisms. Indeed, pre-existing immunity to the endemic coronavirus HCoV-OC43 has been previously reported to associate with survival outcome from COVID-19 infection (*58*). Further investigation into this potential association is needed, given an independent study reporting no correlation of endemic coronavirus immunity and survival after COVID-19 infection (*57*).

The longitudinal characterization of the B cell repertoire of a BNT162b2-vaccinated donor revealed the presence of a cluster of public antibody heavy chain sequences with high similarity to antibody sequences observed in natural infection. These public heavy chain sequences were observed at multiple timepoints after vaccination, with IgM members identified at day 8, and IgG/IgA sequences identified at later timepoints, suggesting antigen-driven evolution of a public mode of SARS-CoV-2 spike recognition. Interestingly, despite the high similarity in heavy chain sequences, antibodies from this public cluster utilized a variety of different light chain V-genes and CDRL3 sequences, suggesting that the immune response to SARS-CoV-2 may utilize a combination of common heavy chain sequence characteristics that are important for antigen recognition, paired with a heterogeneous set of light chain sequences that may allow for additional diversification of the fine epitope specificities of responding B cells. The ability to increase the diversity in B cell responses, while retaining critical antigen interactions, may be an important tool that the human immune system utilizes for counteracting virus evasion mechanisms.

Immunization with BNT162b2 in the donor cohort also resulted in a transient SARS-CoV-2 IgA response. While the polysaccharide pneumococcal vaccine can lead to a persistent IgA vaccine response (*59*), robust IgA responses are not generated by either the intramuscular influenza (*60*) or tetanus-diphtheria-acellular pertussis vaccine (*61*). IgA is critical in the SARS-CoV-2 early neutralizing antibody response (*62*) and was previously observed together with IgG in response to the SARS-CoV-2 mRNA vaccine (*63*). Consistent with the development of protective immunity, antigen-binding B cells were IgG and IgA class-switched and primarily observed among memory B cell and plasmablast subsets. The strong IgA response elicited by the SARS-CoV-2 mRNA vaccines may arise as a consequence of single-stranded RNA directly stimulating B-cells (*64, 65*) or activation of TLR7 on dendritic cells (*66*). Indeed, addition of a single-stranded RNA adjuvant to a traditional influenza vaccine generated mucosal immunity through a robust IgA response and provided more strain cross-protection (*67*).

This study uses unbiased machine learning and sequencing technologies to identify the antigen-specific cells and evolving antibody response to RNA-based vaccination against SARS-CoV-2. The approach allowed isolation of live, virus-specific T cells that will facilitate further studies and development of new therapies. The multi-compartment vaccine response induced by BNT162b2 may have important ramifications, particularly in immunocompromised populations. Reduced humoral responses have been reported to SARS-CoV-2 infection or BNT162b2 in patients with solid organ transplant (*68-70*), cancer (*71*), or immune-mediated inflammatory diseases (*72, 73*). The identification of altered proteins or specific cell populations to serve as biomarkers for successful vaccination may provide important insight to monitor development and continued protection of these vulnerable populations. While limited sample sizes here reduce statistical power for population level correlations and detailed outcomes, the unbiased identification of antigen-specific CD4, CD8, and B cell populations provides important insight into general mechanisms of RNA-based vaccines and the cellular basis for vaccine-induced antibodies and protection from SARS-CoV-2.

## ABBREVIATIONS

BCL6: B-cell lymphoma 6 protein
BCR: B cell receptor
COVID-19: Coronavirus disease 2019
cTfh: Circulating T follicular helper
CyTOF: Cytometry by Time-of-Flight mass spectrometry
CYTOC: Cytochrome C
dsRNA: Double stranded ribonucleic acid
FACS: Fluorescence activated cell sorting
ICOS: Inducible T cell costimulator
LIBRA-seq: Linking B cell receptor to antigen specificity through sequencing
MEM: Marker Enrichment Modeling
mRNA: Messenger ribonucleic acid
SARS-CoV-2: Systemic acute respiratory syndrome coronavirus 2
S protein: Spike protein
TLR: Toll-like receptor
T-REX: Tracking responders expanding
VSV: Vesicular stomatitis virus

## Acknowledgements

We thank members of the Vanderbilt Center for Immunobiology for input and Summer Brown for administrative assistance. This work was supported by Human Immunology Discovery Initiative of the Vanderbilt Center for Immunobiology (JCR, EMW, RHB, JMI), CTSA award No. UL1TR000445 (EMW, JCR), the National Institutes of Health KL2TR002245 (EMW), K12HD043483 (RHB), R01CA226833 (JMI, SMB, CER), U01AI125056 (JMI, SMB), U54CA217450 (JMI), P30CA68485 (VUMC Flow Cytometry Shared Research), R01AI131722-S1 (ISG), R01AI157155 (JEC), R01DK105550 (JCR), R01CA217987 (JCR), R01AI153167 (JCR), R01AI127521 (JSM); HHSN contracts 75N93019C00074 (JEC), DARPA HR0011-18-2-0001 (JEC), Hays Foundation COVID-19 Research Fund (ISG), Fast Grants Mercatus Center of George Mason University (ISG, JEC), the Dolly Parton COVID-19 Research Fund at Vanderbilt (JEC), Welch Foundation F-0003-19620604 (JSM), and William Paul Distinguished Innovator Award of the Lupus Research Alliance (JCR).

## Author Contributions

KJK contributed to conceptualization, data curation, formal analysis, investigation, methodology, visualization, and writing the original draft. EMW contributed to conceptualization, formal analysis, project administration, resources, visualization, and writing the original draft. KV contributed to formal analysis, investigation, methodology, visualization, and writing the original draft. SMB contributed to data curation, formal analysis, visualization, software development, and visualization. CER contributed to investigation and data curation, ARS contributed to conceptualization, data curation, formal analysis, methodology, visualization, software development. NR contributed to data curation, formal analysis, methodology, visualization, software development. NS contributed to data curation. LW contributed to data curation. SCW contributed to data curation. AP contributed to data curation. SS and DD contributed to sample acquisition, processing, and management. CSW, CH, JSM, RHC, and JEC contributed resources. JCR contributed to conceptualization, formal analysis, funding acquisition, methodology, resources, supervision, visualization, and writing the original draft. RHB contributed to conceptualization, data curation, formal analysis, investigation, software, visualization, and writing the original draft. ISG contributed to conceptualization, data curation, funding acquisition, methodology, resources, software, supervision, and visualization. JMI contributed to conceptualization, data curation, formal analysis, funding acquisition, methodology, resources, software, supervision, and visualization.

## Data and materials availability

Materials and data are or will be made available by corresponding authors and deposited in appropriate public databases and materials will be available upon request with appropriate MTA approval. Mass cytometry datasets in this manuscript have been deposited in FlowRepository (http://flowrepository.org/).

## SUPPLEMENTARY FIGURES

**Supplemental Figure 1.**
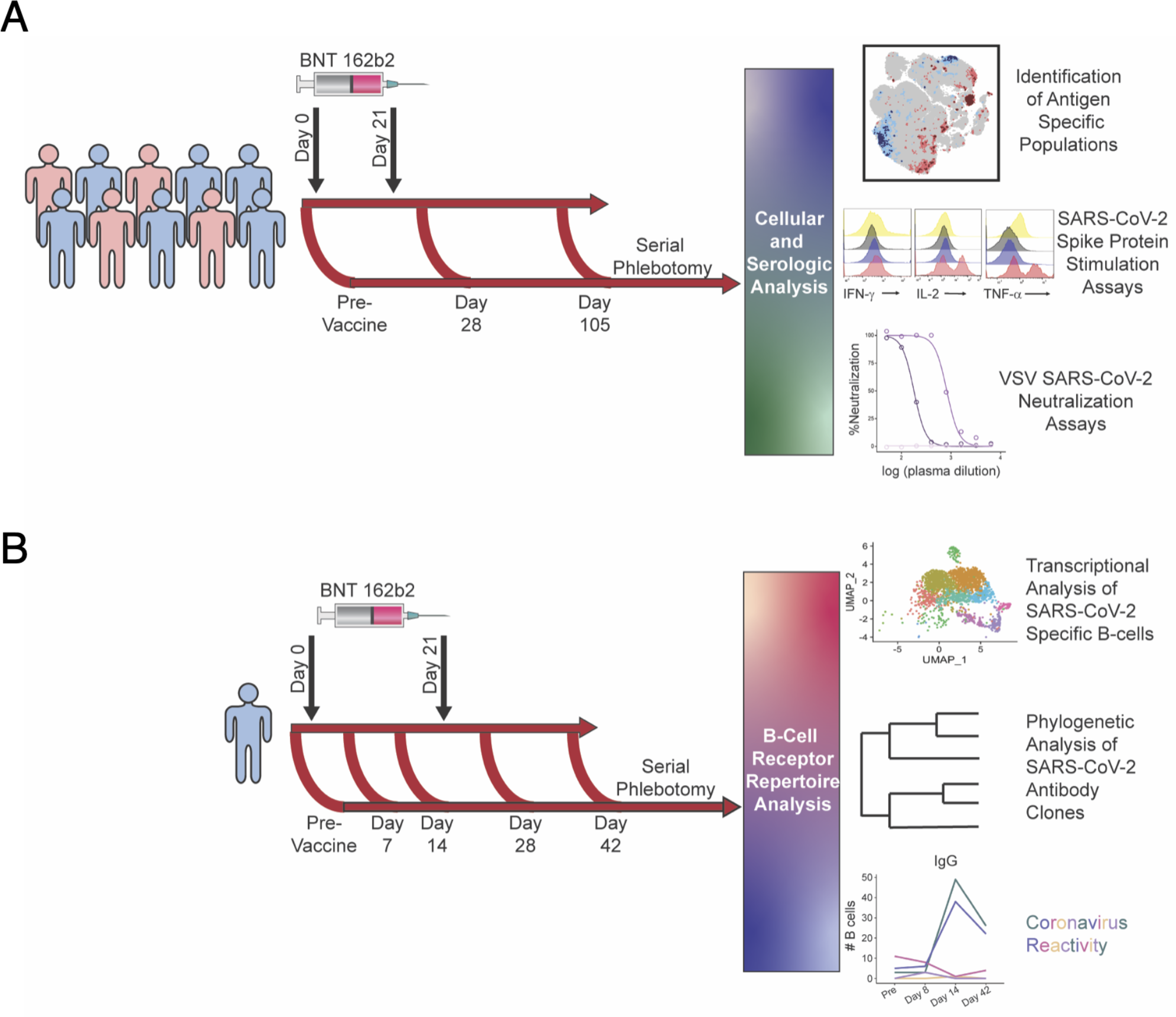
Schematic of vaccination schedule and sample collection. All donors were vaccinated with BNT162b2 on days 0 and 21. **A.** 10 healthy donors underwent serial phlebotomy was performed pre-vaccination (day -3 to 0), on day 28-30, and on day 105-108. PBMCs were isolated at each time point, and citrated plasma was stored when possible. PBMCs from these donors were utilized for CyTOF and *in vitro* stimulation studies. Plasma was used both for SARS-CoV-2 ELISAs and vesicular stomatitis virus pseudoneutralization assays. **B.** A single healthy donor underwent serial phlebotomy pre-vaccination and on days 7, 14, 28, and 42. PBMCs and citrated plasma were isolated at each time point and used for transcriptional analysis of SARS-CoV-2 specific B cells.

**Supplemental Figure 2.**
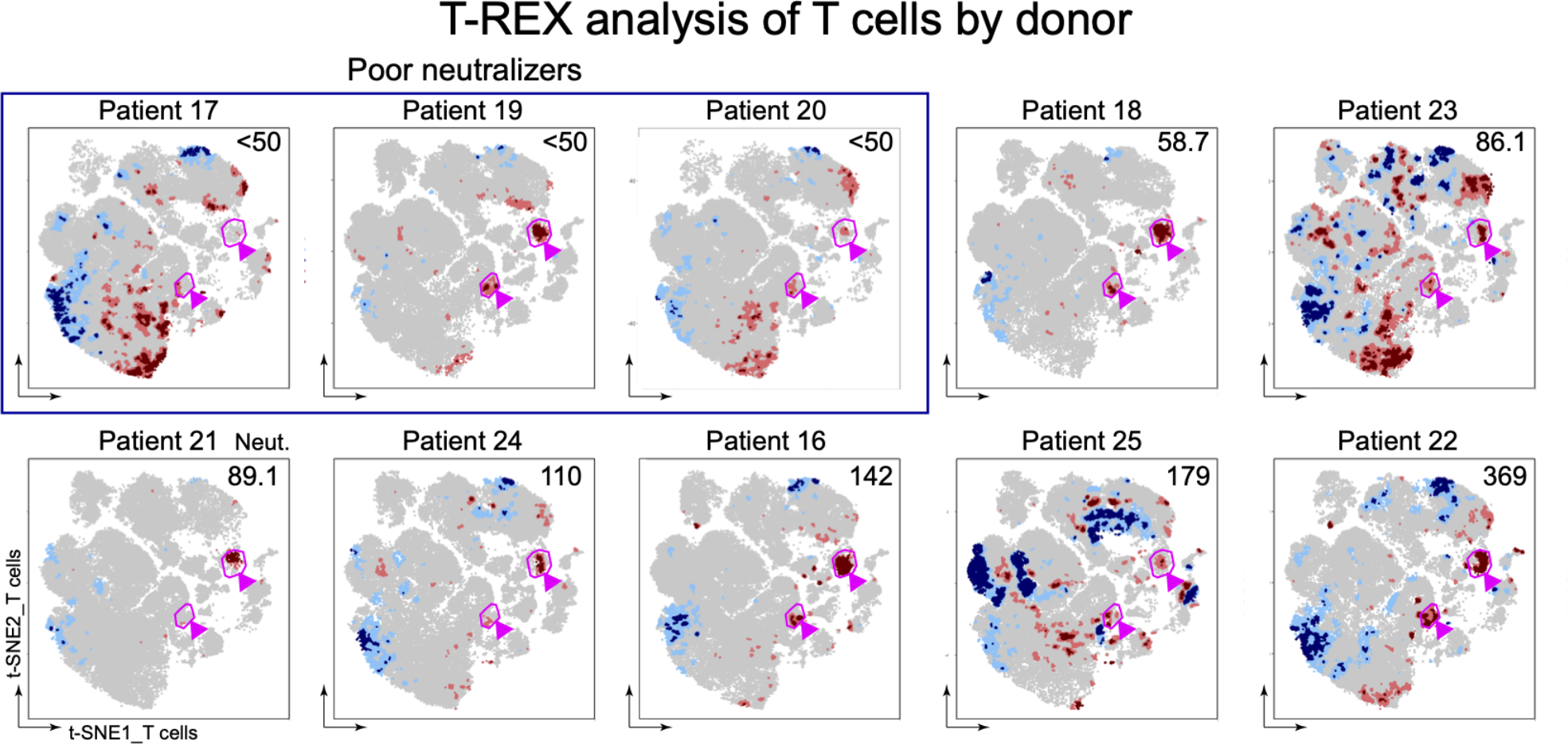
T-REX analysis of T cell populations for individual donors. Individual t-SNE MAPS are shown for each donor. T-REX analysis showing contracting populations (blue) and expanding populations (red) for T cells pre- and post-BNT162b2 vaccination. Pink circles denote the antigen-specific CD8+CD38+ICOS+ T cells (right) and CD4+CD38+ICOS+ T cells (middle) identified in Figure 1. Plasma dilution necessary to achieve 50% neutralization in a SARS-CoV-2 vesicular stomatitis virus pseudonormalization assay is shown in the top right of each map. Donors who failed to achieve 50% neutralization with a 50-fold dilution were deemed poor neutralizers.

**Supplemental Figure 3.**
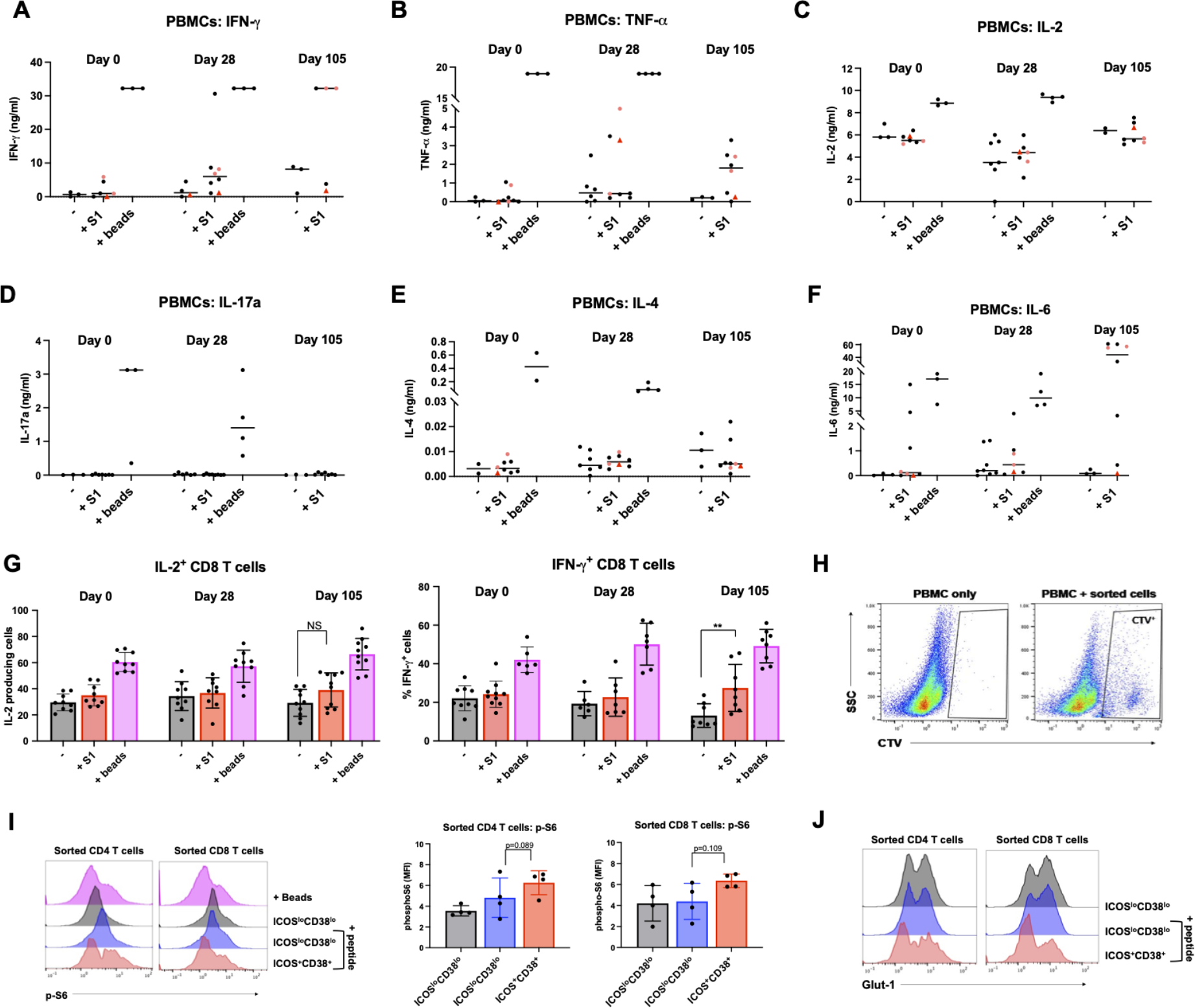
Supporting information for cytokine stimulation assays (Figure 2). PBMCs collected from study participants pre-vaccination (day 0), day 28 post-vaccination, and day 100 post-vaccination were cultured with no stimulation (-), Spike protein (+S1), or polyclonal anti-CD2/3/28 beads. **A-F.** Supernatants were collected from cultures after 4 days and predicted concentrations of each cytokine was determined by Legendplex bead capture assay. **G.** PBMC Cultures were maintained until day 11 post-stimulation and then subjected to flow cytometry to determine percentages of IL-2 and IFN-y producing T cells. **H.** Representative example of CellTraceViolet (CTV)-labeled samples from FACS, reintroduced to Day 0 autologous unlabeled PBMC cultures. **I.** Sorted CTV+ samples were left unstimulated (black) or stimulated with SARS-CoV-2 peptide (blue and red) for 2 days and phospho-S6 was measured by flow cytometry. Significance was determined by paired ANOVA. **J.** Representative Glut-1 staining in sorted samples as in I.

**Supplemental Figure 4.**
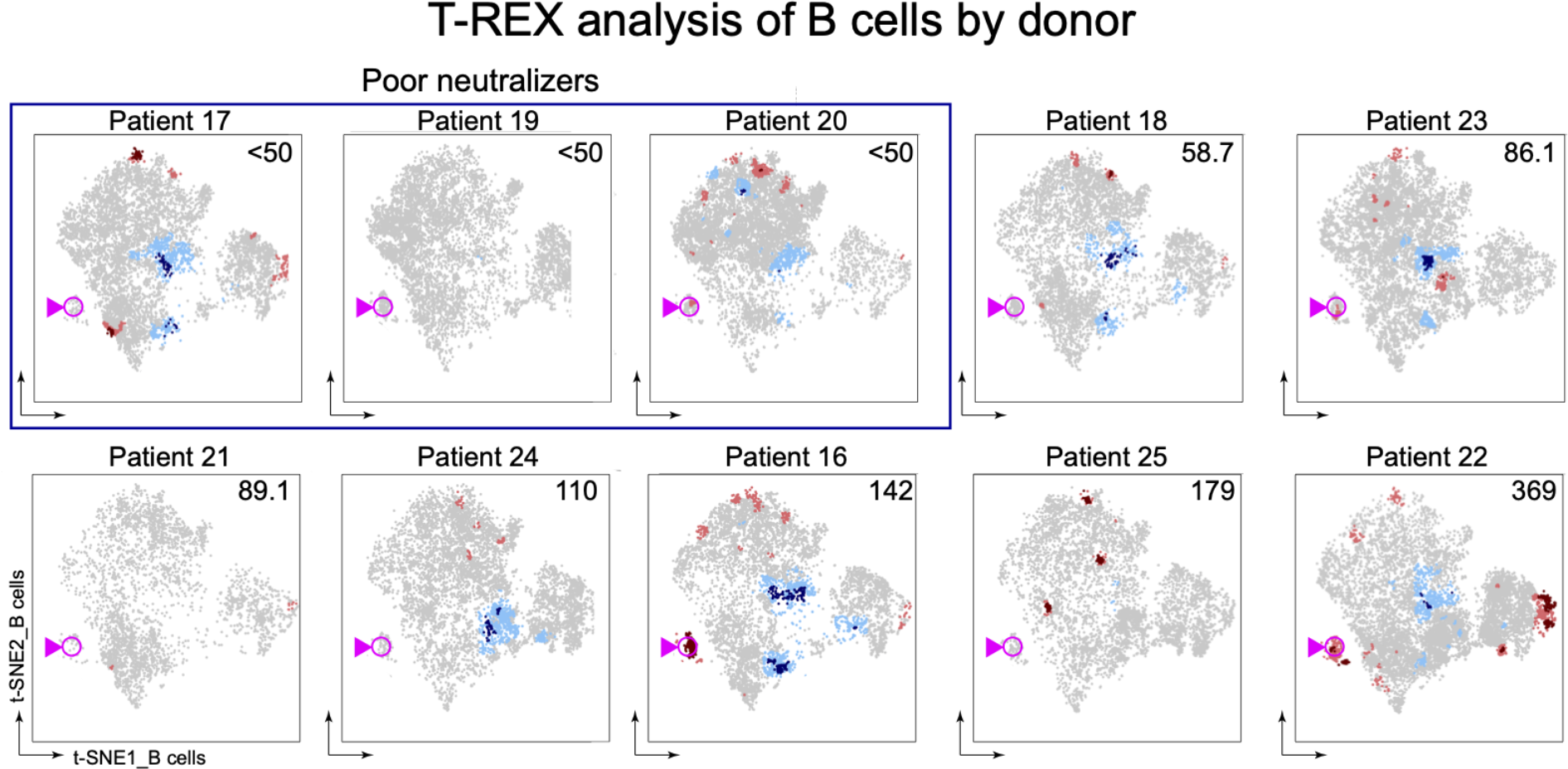
T-REX analysis of B cell populations for individual donors. Individual t-SNE MAPS are shown for each donor. T-REX analysis showing contracting populations (blue) and expanding populations (red) for B cells pre- and post-BNT162b2 vaccination. Pink circles denote the activated plasmablast population. Plasma dilution necessary to achieve 50% neutralization in a SARS-CoV-2 vesicular stomatitis virus pseudonormalization assay is shown in the top right of each map. Donors who failed to achieve 50% neutralization with a 50-fold dilution were deemed poor neutralizers.

**Supplemental Figure 5.**
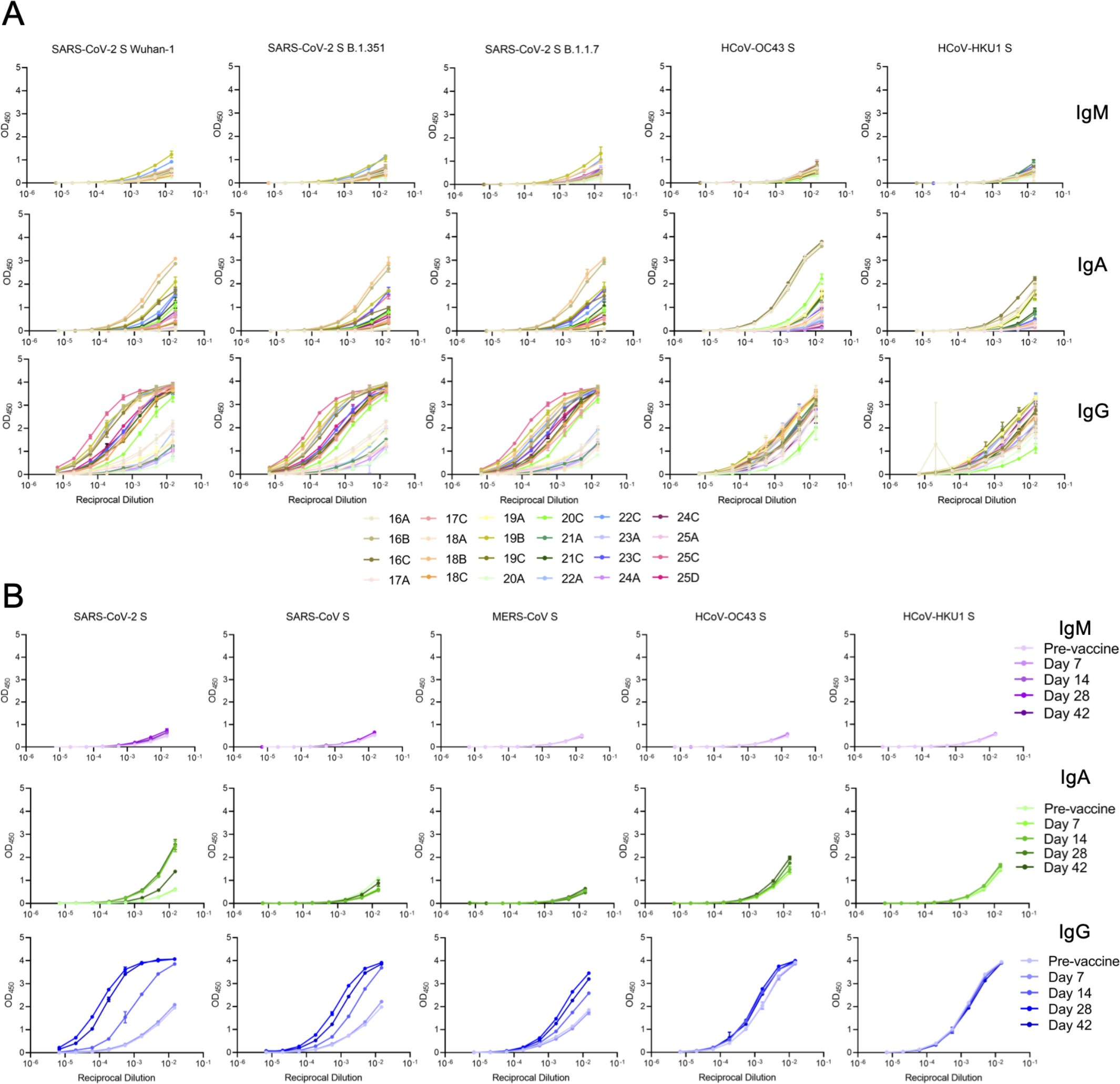
ELISA data for individual donors. **A.** ELISA curves for 10 donor vaccinee plasma for IgM, IgA, and IgG isotypes against spike proteins from SARS-CoV-2, SARS-CoV, MERS-CoV, HCoV-OC43, and HCoV-HKU1. The optical density at 450 nm (y-axis) is plotted as a function of antibody concentration (x-axis). **B.** ELISA curves for longitudinal donor plasma for IgM, IgA, and IgG isotypes against spike proteins from SARS-CoV-2, SARS-CoV, MERS-CoV, HCoV-OC43, and HCoV-HKU1. The optical density at 450 nm (y-axis) is plotted as a function of antibody concentration (x-axis).

**Supplemental Figure 6.**
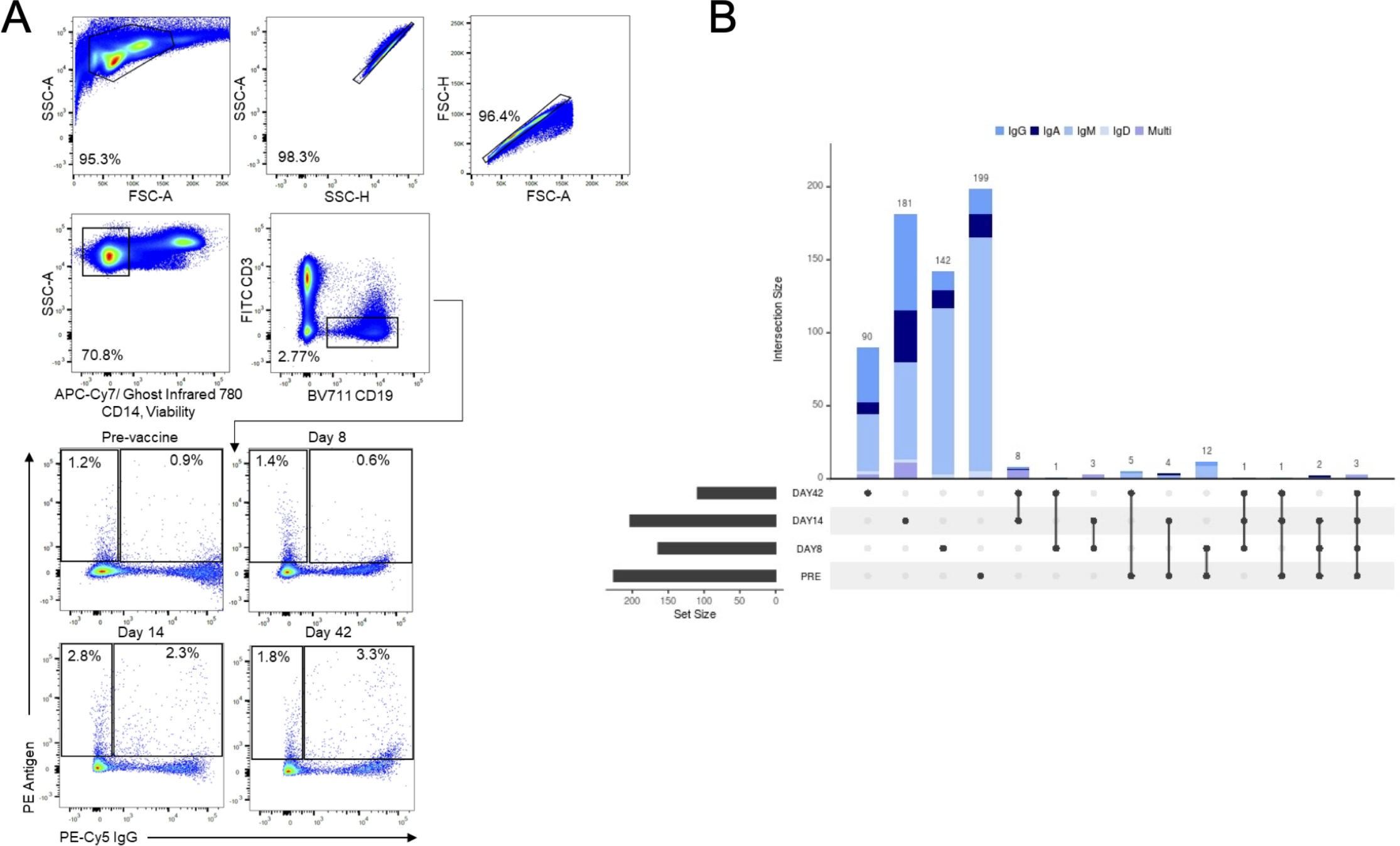
Flow cytometry enrichment of coronavirus antigen-binding B cells. **A.** Gating scheme for fluorescence activated cell sorting (FACS) of PBMCs. Cells were stained with viability dye Ghost Infrared 780, CD14-APC-Cy7, CD3-FITC, BV711-CD19, and IgG-PE-Cy5. Antigen positive B cells were detected by streptavidin-PE binding to biotinylated DNA-oligo tagged antigens. Gates are drawn based on parameters set during cell sorting and percentages are from sort are listed in each plot. The gate for enrichment of CD19+ cells is shown for the pre-vaccine sample; Day 8, Day 14, and Day 42 samples all followed this sorting scheme. **B.** Representation of unique and shared clonotypes between different timepoints. For each individual and combination of timepoints, the number of clonotypes is displayed as a vertical bar graph. Shared clonotypes between timepoints is displayed by filled circles, showing which timepoints are part of a given shared cluster/combination where each combination is mutually exclusive. For each timepoint, the total number unique and shared clonotypes is indicated as a horizontal bar at the bottom left of the panel. Isotypes of clonotypes are represented in different colors in the vertical bar graphs. Clusters with multiple isotypes is shown as “Multi.”

**Supplemental Figure 7.**
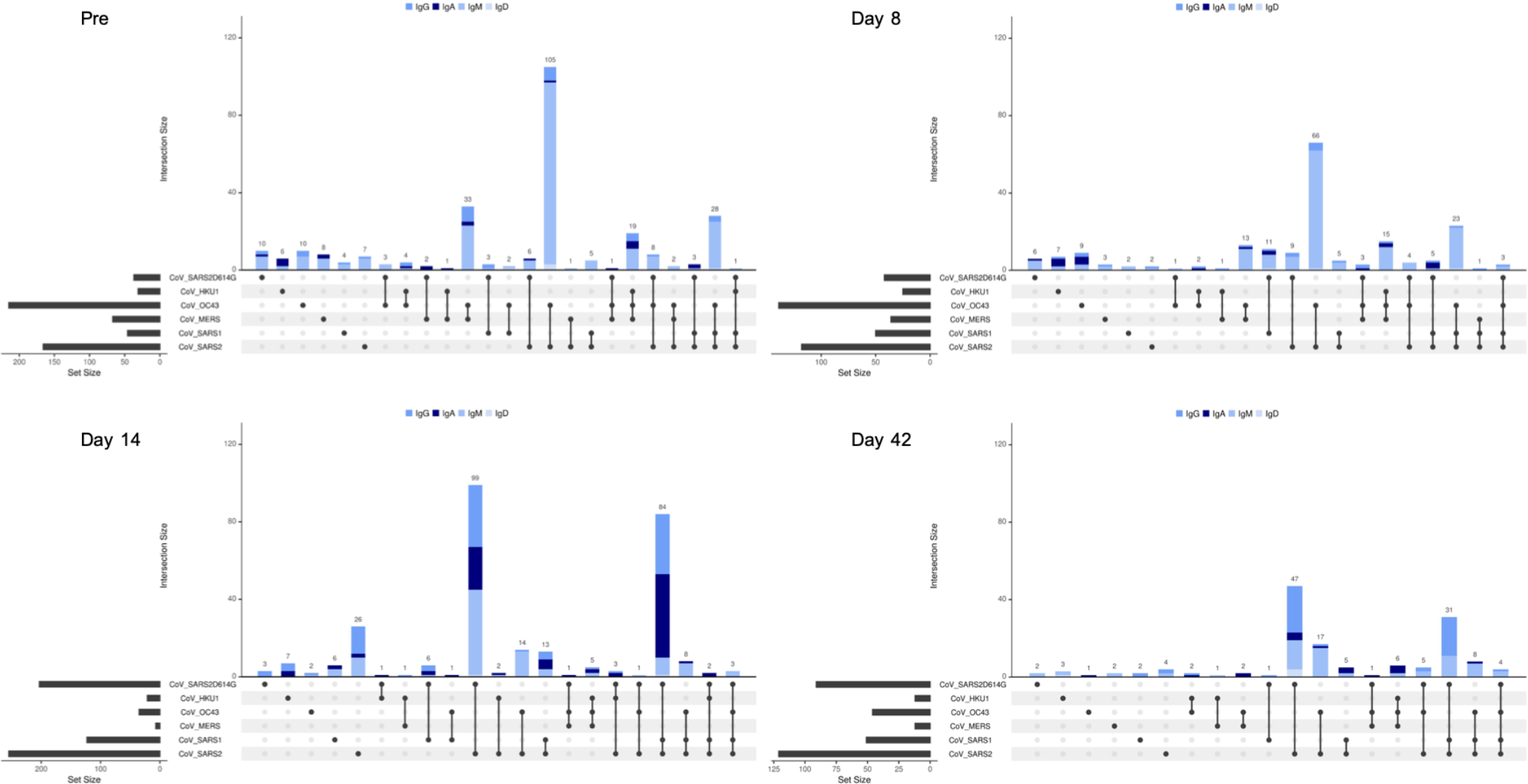
B cell clones with cross-reactive LIBRA-seq scores over time. For each combination of CoV antigens, the number of B cells with high LIBRA-seq scores (≥1) is displayed as a bar graph. The combination of antigens is displayed by filled circles, showing which antigens are part of a given combination. Each combination is mutually exclusive. The number of B cells with high LIBRA-seq scores for each antigen is indicated as a horizontal bar at the bottom left of the panel. Isotypes of select B cells are depicted as more intense colors of blue beginning at IgD and following with IgM, IgG, and IgA.

**Supplemental Figure 8.**
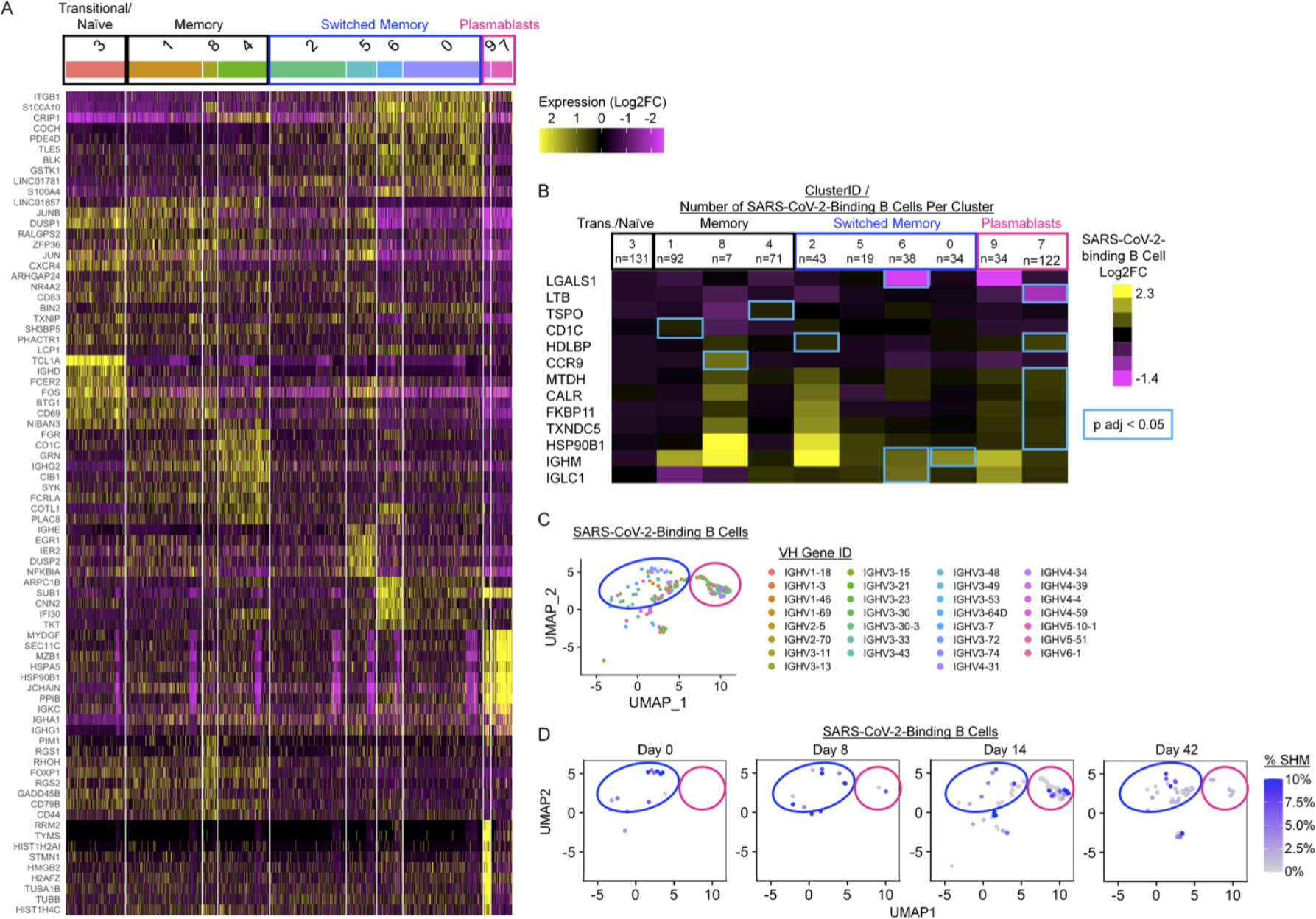
Single-cell RNAseq analysis identifies distinct clusters of memory and plasmablast subsets among B cells enriched for coronavirus antigen-binding by LIBRAseq. **A.** The top 10 differentially expressed genes were identified by Seurat for each cluster identified as in Fig. 5. Heatmap depicts relative expression for each gene, columns are individual cells within each cluster. **B-D.** SARS-CoV-2-binding B cells were identified by LIBRAseq as in Fig. 5. Circles depict switched memory (blue) and plasmablast clusters (pink). **B.**SARS-CoV-2-binding B cells were compared to non-SARS-CoV-2-binding B cells within each cluster identified in panel A. Log2 fold change is shown for differentially-expressed genes that were significantly different between these two groups in at least one cluster (p adj < 0.05, blue rectangles). **C.** VH gene identity is shown among all SARS-CoV-2-binding B cells.**D.** Somatic hypermutation is shown among SARS-CoV-2-binding B cells identified at each time point following vaccination with BNT162b2.

**Supplemental Figure 9.**
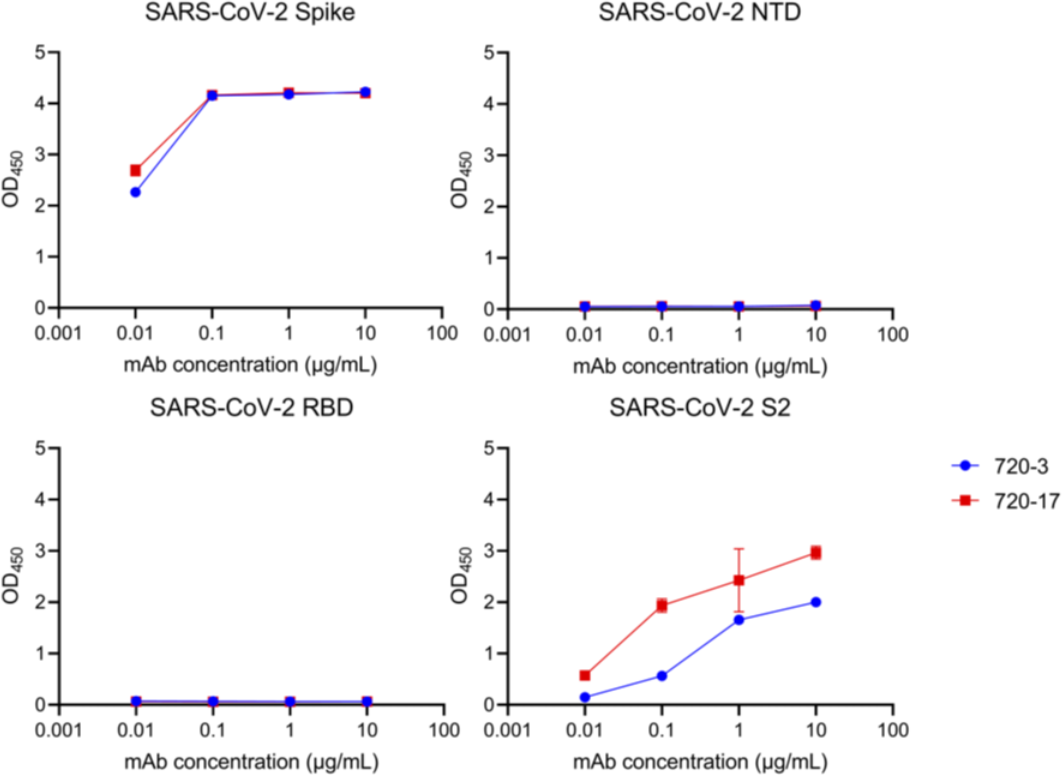
Domain mapping of recombinant monoclonal antibodies. ELISA data for the SARS-CoV-2 spike and the truncated SARS-CoV-2 NTD, RBD, and S2 domains. Optical density at 450 nm (y-axis) is depicted as a function of antibody concentration (x-axis).

**Supplemental Table 1.**
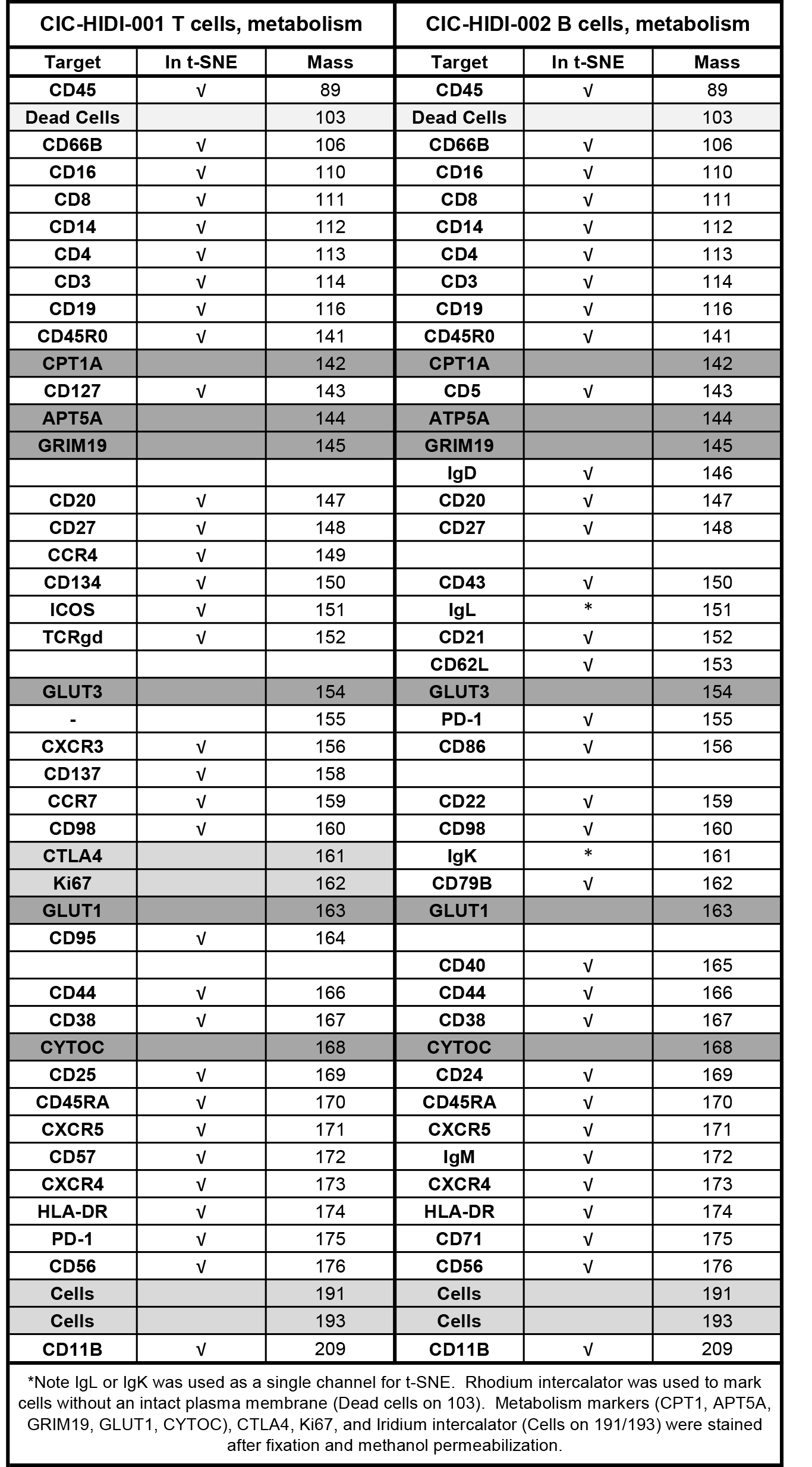
Mass cytometry antibody panels and clones.

## MATERIALS AND METHODS

### Human Subjects Information

IRB approval was obtained (VUMC 191562). Informed consent was obtained, and a baseline health questionnaire was also completed. Simple phlebotomy was performed either pre-vaccine (day 0), day 28-30, and day 95-100 OR pre-vaccine and days 8, 14, and 42 after initial BNT162b2 vaccination using sodium citrate mononuclear cell preparation (CPT) tubes. All participants received two doses of vaccine 21 days apart.

### PBMC and Plasma isolation

CPT tubes were spun at 1600 RCF for 20 minutes. The plasma layer was carefully removed, transferred to a conical vial, spun at 600g for 10 minutes, and the supernatant transferred to microtubes in 1 mL aliquots. Plasma was stored at -80°C until further use. Buffy coat was divided amongst two clean conical tubes. CPT tubes were rinsed with 1 mL PBS (Gibco), and the total volume in the conical was increased to 15cc. Cells were pelleted at 600g for 10 minutes. Cell pellets were combined and washed with 10cc PBS, and then cells were pelleted again at 600g for 10 minutes. PBS was discarded, and the pellet was re-suspended in 3mL ACK buffer (Gibco) for 5 minutes. 10 mL of PBS was added to the ACK cell suspension, and cells were pelleted for 10 minutes at 600g. Cells were resuspended in PBS, strained through a cell strainer (Falcon) and counted using an ACT Diff hematology analyzer (Beckman Coulter). Cells were pelleted by centrifugation at 600g for 10 minutes and resuspended in heat-inactivated FBS (Gibco) containing 10% DMSO (Sigma-Aldrich) at a concentration of 5 million cells/mL in cryovials (Nalgene). Cryovials were frozen overnight to -80°C using Mr. Frosty freezing containers (Nalgene) and then transferred to liquid nitrogen for long-term storage.

### Mass Cytometry

#### PBMC Staining

Metal-tagged antibodies were purchased from Fluidigm. Cell labeling and mass cytometry analysis were performed as previously described (*1, 2*). Briefly, cells were incubated with a viability reagent (Cell ID Intercalator-Rh; Fluidigm), per the product literature. Then cells were washed in PBS without calcium or magnesium (Gibco, Thermo Fisher Scientific) containing 1% BSA (Thermo Fisher Scientific) and stained in 50 μL PBS and BSA 1%–containing antibody cocktail for extracellular targets. Cells were stained for 30 minutes at room temperature using the antibodies listed in Supplemental Table 3. Cells were washed in PBS and BSA 1% and then fixed with 1.6% paraformaldehyde (Electron Microscopy Sciences). Cells were washed once in PBS and permeabilized by resuspension in ice-cold methanol. After incubation overnight at −20°C, cells were washed with PBS and BSA 1% and stained in 50 μL PBS and BSA 1%– containing antibody cocktail for intracellular targets. Cells were washed in PBS and BSA 1%, then washed with PBS and stained with an iridium DNA intercalator (Fluidigm) for 20 minutes at room temperature. Finally, cells were washed with PBS and with diH2O before being resuspended in 1× EQ Four Element Calibration Beads (Fluidigm) and collected on a Helios mass cytometer (Fluidigm) at the Vanderbilt Flow Cytometry Shared Resource Center. Events were normalized as previously described (*3*).

#### Data analysis

After normalization, CYTOF data were scaled with an arcsinh transformation, with an appropriate cofactor set for each channel following standard procedures for fluorescence and mass cytometry data (*4*). Data were then manually gated for removal of atypical events (*5*). After quality control gating, a UMAP analysis was performed on the cleaned-up samples using the surface markers in each panel. Metabolic markers, ki67, iridium, and rhodium were not used to create the UMAP. The resulting common, 2-dimensional embedding of the data was used for visualization and selection of either CD3+ T cells (from T cell panel data) or B cells (from B cell panel data) for further downstream analysis. A common t-SNE analysis was done on all 10 donor samples using the same markers used to create the UMAP on either the CD3+ T cells or B cells extracted from their respective UMAP. After the t-SNE, equally sampling was done on each donor pair. The 10 donor samples were then combined for a T-REX comparison of day 0 and day 28 data (*6*). MEM was used to quantify enriched features in each region of significant change (*7*). Comparisons of population frequencies pre- and post-vaccination as well as correlations between post-vaccine cell frequencies and IgG titers were done in GraphPad Prism version 9.0. Populations were compared using Mann-Whitney U tests. Statistical correlations were determined using Pearson correlations. P values less than 0.05 were considered statistically significant.

### *In vitro* characterization of CD38+ICOS+ Cells

#### Quantification of CD38+ICOS+ CD4 and CD8 T cells by flow cytometry

To quantify CD38+ICOS+ CD4 and CD8 T cells, PBMCs were first resuspended with Human TruStain Fcx (Biolegend) for 10 minutes at room temperature and then stained with the following antibodies in FACS buffer (PBS+ 2% fetal bovine serum): CD8a e450 (Invitrogen 48-0086-42, 1:200) ICOS BV605 (Biolegend 313538, 1:50), CCR7 PE (Biolegend 353204, 1:200), CD38 PerCP (Biolegend 303520, 1:100), CD4 PECy7 (Biolegend 357410, 1:100), and CD3 APCCy7 (Biolegend 300318, 1:200). Cells were analyzed on a Miltenyi MACSQuant16 Analyzer with single-stain control PBMC samples used for compensation conducted in FlowJo v10.6.2.

#### Stimulation of ICOS+CD38+ and ICOS^lo^CD38^lo^ populations with SARS-CoV-2 S protein

The same staining procedure was used for FACS of ICOS+CD38+ and ICOSloCD38lo populations on a BD FACSAria III instrument in the Vanderbilt University Medical Center Flow Cytometry Shared Resource Core. Sorted cells were stained with 1uM CellTraceViolet (CTV) (ThermoFisher) for 20 minutes and then cultured with autologous unlabeled PBMC samples from day 0 (pre-vaccination) samples. Before adding CTV-labeled cells to PBMCs for activation, PBMCs were CD3-depleted by positive selection (CD3 Microbeads, Miltenyi Biotec) and purity of CD3 depletions were assessed by flow cytometry. Cells were cultured in Human Plasma-like Media (HPLM) (*8*) + 1% pen/strep + 5% dialyzed serum (Sigma Aldrich). For SARS-CoV-2 peptide stimulation, cultures were treated with Peptivator SARS-CoV2 Prot_S peptide pool (Miltenyi Biotec #130-126-700) in a 48-well plate format for 2 days.

For cytokine staining, cells were either stimulated with PMA (1ug/ml) and ionomycin (750ng/ml) for 5 hours, or restimulated on day 11 in 96-well plates with 2.5ug/ml recombinant SARS-CoV-S protein S1 (Biolegend 792906, carrier-free) for 8h in the presence of 1ug/ml Golgiplug and 0.7ug/ml Golgistop. Peptivator-stimulated cultures were treated with Golgiplug/Golgistop overnight after 2 days of activation. Cells were surface stained in FACS buffer, fixed with 1.5% paraformaldehyde for 10 minutes, and permeabilized with methanol for 20 minutes on ice. Additional intracellular antibodies were: TNF-a AF488 (Biolegend 502915, 1:100), IFN-g APC (Invitrogen 17-7319-82, 1:100) IL-2 AlexaFluor 700 (Biolegend 500320, 1:150), granzyme b FITC (Biolegend 515403, 1:100), phospho-S6 APC Ser235/236 (Invitrogen 17-9007-42, 1:80), Bcl-6 FITC (Biolegend 358513, 1:100), and Glut-1 AlexaFluor 647 (Abcam ab115730, 1:300). Cytokines measured from PBMC supernatants were collected after 4 days of incubation with S protein and concentrations were predicted using a standard curve in the LEGENDplex assay (Miltenyi Biotec 741028).

#### Data Analysis

Group comparisons were performed using GraphPad Prism version 9.0. Populations were compared using Mann-Whitney U tests. P values less than 0.05 were considered statistically significant.

### B Cell Receptor Repertoire and Serologic Analysis

#### Single cell profiling of antigen specific B cells

##### Recombinant expression and purification of coronavirus antigens

Plasmids encoding residues 1–1208 of the SARS-CoV-2 spike with a mutated S1/S2 cleavage site, proline substitutions at positions 817, 892, 899, 942, 986 and 987, and a C-terminal T4-fibritin trimerization motif, an 8x HisTag, and a TwinStrepTag (SARS-CoV-2 S HP); 1–1208 of the SARS-CoV-2 spike with a mutated S1/S2 cleavage site, proline substitutions at positions 817, 892, 899, 942, 986 and 987, as well as mutations L18F, D80A, L242-244L del, R246I, K417N, E484K, N501Y, and a C-terminal T4-fibritin trimerization motif, an 8x HisTag, and a TwinStrepTag (SARS-CoV-2 S HP Beta); 1–1208 of the SARS-CoV-2 spike with a mutated S1/S2 cleavage site, proline substitutions at positions 817, 892, 899, 942, 986 and 987, as well as mutations 69-70del, Y144del, N501Y, A570D, P681H, and a C-terminal T4-fibritin trimerization motif, an 8x HisTag, and a TwinStrepTag (SARS-CoV-2 S HP Alpha); residues 1-1190 of the SARS-CoV spike with proline substitutions at positions 968 and 969, and a C-terminal T4-fibritin trimerization motif, an 8x HisTag, and a TwinStrepTag (SARS-CoV S-2P); residues 1-1291 of the MERS-CoV spike with a mutated S1/S2 cleavage site, proline substitutions at positions 1060 and 1061, and a C-terminal T4-fibritin trimerization motif, an AviTag, an 8x HisTag, and a TwinStrepTag (MERS-CoV S-2P Avi); residues 1-1277 of the HCoV-HKU1 spike with a mutated S1/S2 cleavage site, proline substitutions at positions 1067 and 1068, and a C-terminal T4-fibritin trimerization motif, an 8x HisTag, and a TwinStrepTag (HCoV-HKU1 S-2P); residues 1-1278 of the HCoV-OC43 spike with proline substitutions at positions 1070 and 1071, and a C-terminal T4-fibritin trimerization motif, an 8x HisTag, and a TwinStrepTag (HCoV-OC43 S-2P); were transiently transfected into FreeStyle293F cells (Thermo Fisher) using polyethylenimine. For all antigens with the exception of SARS-CoV-2 S HP, cells were treated with 1 µM kifunensine to ensure uniform glycosylation three hours post-transfection. Transfected supernatants were harvested after five days of expression. SARS-CoV-2 S HP, SARS-CoV S-2P, MERS-CoV S-2P, HCoV-HKU1 S-2P and HCoV-OC43 S-2P were purified using StrepTrap columns. SARS-CoV-2 S HP, SARS-CoV-2 S HP Beta, SARS-CoV-2 S HP Alpha, SARS-CoV S-2P, MERS-CoV S-2P, HCoV-HKU1 S-2P and HCoV-OC43 S-2P were purified over a Superose6 Increase column (GE Life Sciences).

##### Recombinant expression and purification of ZM197 Env and NC99 Hemagglutinin

Recombinant, soluble HIV-1 gp140 SOSIP trimer from strain ZM197 (clade) containing an AviTag and recombinant NC99 HA protein consisting of the HA ectodomain with a point mutation at the sialic acid-binding site (Y98F) to abolish non-specific interactions, a T4 fibritin foldon trimerization domain, AviTag, and hexahistidine-tag, were expressed in Expi 293F cells using polyethylenimine transfection reagent and cultured. FreeStyle F17 expression medium supplemented with pluronic acid and glutamine was used. The cells were cultured at 37°C with 8% CO2 saturation and shaking. After 5-7 days, cultures were centrifuged and supernatant was filtered and run over an affinity column of agarose bound Galanthus nivalis lectin. The column was washed with PBS and antigens were eluted with 30 mL of 1M methyl-a-D-mannopyranoside. Protein elutions were buffer exchanged into PBS, concentrated, and run on a Superdex 200 Increase 10/300 GL Sizing column on the AKTA FPLC system.

##### Biotinylation of antigens

Constructs containing an Avi-tag (ZM197 Env and HA NC99) were biotinylated using the site-specific biotinylation kit according to manufacturer instructions (Avidity LLC.) All other antigens not containing an Avi-tag were non-specifically biotinylated using the EZ-Link Sulfo-NHS-Biotin kit at a 50:1 biotin:protein molar ratio.

##### DNA-barcoding of antigens

We used oligos that possess 15 bp antigen barcode, a sequence capable of annealing to the template switch oligo that is part of the 10X bead-delivered oligos, and contain truncated TruSeq small RNA read 1 sequences in the following structure: 5’-CCTTGGCACCCGAGAATTCCANNNNNNNNNNNNNCCCATATAAGA*A*A-3’, where Ns represent the antigen barcode. We used the following antigen barcodes: GCAGCGTATAAGTCA (SARS-CoV-2 S HP), GACAAGTGATCTGCA (SARS-CoV-2 S HP D614G), GCTCCTTTACACGTA (SARS-CoV S), GGTAGCCCTAGAGTA (MERS-CoV S), TGTGTATTCCCTTGT (HCoV-HKU1 S), AGACTAATAGCTGAC (HCoV-OC43 S), TCATTTCCTCCGATT (ZM197 EnV), CTTCACTCTGTCAGG (HA NC99), Oligos were ordered from IDT with a 5’ amino modification and HPLC purified.

For each antigen, a unique DNA barcode was directly conjugated to the antigen itself. In particular, 5’amino-oligonucleotides were conjugated directly to each antigen using the Solulink Protein-Oligonucleotide Conjugation Kit (TriLink cat no. S-9011) according to manufacturer’s instructions. Briefly, the oligo and protein were desalted, and then the amino-oligo was modified with the 4FB crosslinker, and the biotinylated antigen protein was modified with S-HyNic. Then, the 4FB-oligo and the HyNic-antigen were mixed together. This causes a stable bond to form between the protein and the oligonucleotide. The concentration of the antigen-oligo conjugates was determined by a BCA assay, and the HyNic molar substitution ratio of the antigen-oligo conjugates was analyzed using the NanoDrop according to the Solulink protocol guidelines.

AKTA FPLC was used to remove excess oligonucleotide from the protein-oligo conjugates, which were also verified using SDS-PAGE with a silver stain. Antigen-oligo conjugates were also used in flow cytometry titration experiments.

##### Antigen specific B cell sorting

Cells were stained and mixed with DNA-barcoded antigens and other antibodies, and then sorted using fluorescence activated cell sorting (FACS). First, cells were counted and viability was assessed using Trypan Blue. Then, cells were washed three times with DPBS supplemented with 0.1% Bovine serum albumin (BSA). Cells were resuspended in DPBS-BSA and stained with cell markers including viability dye (Ghost Red 780), CD14-APC-Cy7, CD3-FITC, CD19-BV711, and IgG-PE-Cy5. Additionally, antigen-oligo conjugates were added to the stain (1 µg of every antigen except for HA NC99 and HCoV-HKU1 S which were added at 0.1 µg). After staining in the dark for 30 minutes at room temperature, cells were washed three times with DPBS-BSA at 300 g for five minutes. Cells were then incubated for 15 minutes at room temperature with Streptavidin-PE to label cells with bound antigen. Cells were washed three times with DPBS-BSA, resuspended in DPBS, and sorted by FACS. Antigen positive cells were bulk sorted and delivered to the Vanderbilt Technologies for Advanced Genomics (VANTAGE) sequencing core at an appropriate target concentration for 10X Genomics library preparation and subsequent sequencing. FACS data were analyzed using FlowJo.

##### Sample preparation, library preparation, and sequencing

Single-cell suspensions were loaded onto the Chromium Controller microfluidics device (10X Genomics) and processed using the B-cell Single Cell V(D)J solution according to manufacturer’s suggestions for a target capture of 10,000 B cells per 1/8 10X cassette, with minor modifications in order to intercept, amplify and purify the antigen barcode libraries as previously described (*9*).

##### Sequence processing and bioinformatic analysis

We utilized our previously described pipeline to use paired-end FASTQ files of oligo libraries as input, process and annotate reads for cell barcode, UMI, and antigen barcode, and generate a cell barcode - antigen barcode UMI count matrix (*10*). BCR contigs were processed using Cell Ranger (10X Genomics) using GRCh38 as reference. Antigen barcode libraries were also processed using Cell Ranger (10X Genomics). The overlapping cell barcodes between the two libraries were used as the basis of the subsequent analysis. We removed cell barcodes that had only non-functional heavy chain sequences as well as cells with multiple functional heavy chain sequences. Additionally, we aligned the BCR contigs (filtered_contigs.fasta file output by Cell Ranger, 10X Genomics) to IMGT reference genes using HighV-Quest (*11*). The output of HighV-Quest was parsed using Change-O (*12*). and merged with an antigen barcode UMI count matrix. Finally, we determined the LIBRA-seq score for each antigen in the library for every cell as previously described (*9*).

##### LIBRA-seq data quality control filtering

Cells were filtered based on multiple criteria for further analysis. Cells were only included if the sum of all antigen UMI counts for a particular cell barcode was greater than 4. All cells that met these criteria from pre, day 8, day 14, and day 42 time points were combined. Then, we removed cells from the dataset that had multiple heavy chains or multiple light chains associated with a single cell barcode. Then, LIBRA-seq scores were generated (8). Briefly, a pseudocount of 1 was added to each antigen UMI count, and then centered-log ratios (CLR) were calculated for each antigen UMI count for each cell. Then, an antigen-wise z-score transformation was applied. After performing the LIBRA-seq score calculation, cells were filtered out if they fulfilled any of the following criteria: (1) Max UMI among all antigens less than or equal to 30, (2) ZM197 UMI counts greater than or equal to 30, (3) HA UMI counts greater than or equal to 30, (4) MERS UMI counts greater than 10 times the max UMI of other CoV in that cell, or (5) 10 times ZM197 UMI counts greater than the max UMI of non-MERS CoV in that cell. **Supplemental Figure 6B** and **Figure 7B** were generated using the pre-filtered data. **Supplemental Figure 7** and **Figure 5C-E** used post-filtered data.

##### Antibody sequence clustering

Using pre-filtered data, cells from multiple timepoints were combined together to identity highly similar antibody sequences between timepoints. Single-linkage clustering was performed using Change-O (*12*) with the criteria of same VH- and JH-gene usage, same junction and CDRH3 length and 80% CDRH3 nucleotide sequence identity.

##### Phylogenetic trees

For the selected cluster 720, phylogenetic analysis was performed to understand the heavy chain sequence similarities. The sequences from cluster 720 used IGHV3-30-3 and IGHJ4 genes and included 18 members from 3 different post-vaccination timepoints. The cluster 720 heavy chain sequences were compared to sequences from the Coronavirus antibody database (CoV-AbDab) to identify antibodies with high sequence similarity (based on same VH-and JH-gene usage and 70% CDRH3 amino acid sequence identity) to at least one member of cluster 720. This resulted in the identification of 14 antibody sequences that fulfilled the similarity criteria with the cluster 720 sequences. To generate a phylogenetic tree of these public sequences, the tree was rooted on a putative germline sequence that was generated using the IGHV3-30-3 gene, a consensus CDRH3 sequence based on the cluster 720 members with 100% germline gene identity, and the IGHJ4 J-gene for framework 4. The tree was visualized using the Dendroscope tool (*13*).

##### Plasma Serology ELISAs

100 µL of antigen was plated at a concentration of 500 ng/µL in PBS overnight at 4°C. The following day plates are washed with 1X PBS and 0.1% Tween-20 (PBS-T) and then blocked with 5% non-fat dry milk (NFDM). After an hour blocking at 1 hour at RT, plates are washed 3X with PBS-T and plasma is diluted in 1% NFDM PBS-T at a top concentration of 1:67 followed by 7 3-fold dilutions. The plates were incubated at RT for 1 hour and then washed three times in PBST. The secondary antibody was added in 1% NFDM in PBS-T to the plates, which were incubated for one hour at RT. Plates were washed three times with PBS-T and then developed by adding TMB substrate to each well. The plates were incubated at room temperature for ten minutes, and then 1N sulfuric acid was added to stop the reaction. Plates were read at 450 nm.

##### Plasma VSV SARS-CoV-2 Neutralization Assay

In brief, 100uL of plasma samples are heat inactivated at 570C for 1hr and starting at 1:25 dilution eight 2-fold serial dilutions were made with DMEM supplemented with 2% FBS. To determine neutralizing activity of plasma/serum, we used real-time cell analysis (RTCA) assay on an xCELLigence RTCA MP Analyzer (ACEA Biosciences Inc.) that measures virus-induced <u>cytopathic effect</u> (CPE). Briefly, 50 μL of cell culture medium (DMEM supplemented with 2% FBS) was added to each well of a 96-well E-plate using a ViaFlo384 liquid handler (Integra Biosciences) to obtain background reading. A suspension of 18,000 Vero cells in 50 μL of cell culture medium was seeded in each well, and the plate was placed on the analyzer. Measurements were taken automatically every 15 min, and the sensograms were visualized using RTCA software version 2.1.0 (ACEA Biosciences Inc). VSV-SARS-CoV-2 (0.01 MOI, ∼120 PFU per well) was mixed 1:1 with a dilution of plasma/serum in a total volume of 100 μL using DMEM supplemented with 2% FBS as a diluent and incubated for 1 h at 37°C in 5% CO2. At 16 h after seeding the cells, the virus-plasma/serum mixtures were added in replicates to the cells in 96-well E-plates. Triplicate wells containing virus only (maximal CPE in the absence of mAb) and wells containing only Vero cells in medium (no-CPE wells) were included as controls. Plates were measured continuously (every 15 min) for 48 h to assess virus neutralization. Normalized cellular index (CI) values at the endpoint (48 h after incubation with the virus) were determined using the RTCA software version 2.1.0 (ACEA Biosciences Inc.). Results are expressed as percent neutralization in a presence of respective plasma/serum relative to control wells with no CPE minus CI values from control wells with maximum CPE. RTCA IC50 values were determined by nonlinear regression analysis using Prism software.

##### Single-Cell RNA-Seq analysis

Single-cell analysis was performed using Seurat v4.0.0 (*14*). Cells with fewer than 200 RNA features that contained greater than 10% mitochondrial genes were removed. Immunoglobulin VH, Vκ, and Vλ genes were removed prior to UMAP clustering of RNAseq data to prevent them from driving transcriptionally-defined clusters. LIBRA-seq scores were used to identify SARS-CoV-2-binding B cells. B cell subset identities were assigned to clusters based on transcriptional profiles that were consistent with other studies defining these populations. The Seurat FindMarkers function, which uses a non-parametric Wilcoxon rank sum test, was used to identify differentially expressed genes between SARS-CoV-2-binding B cells and non-SARS-CoV-2-binding B cells within each transcriptionally-defined cluster. VDJ data were processed via CellRanger, IMGT/HighV-QUEST, and CHANGE-O, as outlined above to assign V genes, isotypes, and calculate percent VH somatic hypermutation.

## REFERENCES

1. E. Dong, H. Du, L. Gardner, An interactive web-based dashboard to track COVID-19 in real time. Lancet Infect Dis 20, 533–534 (2020).

2. J. Schulte-Schrepping et al., Severe COVID-19 Is Marked by a Dysregulated Myeloid Cell Compartment. Cell 182, 1419–1440 e1423 (2020).

3. D. Mathew et al., Deep immune profiling of COVID-19 patients reveals distinct immunotypes with therapeutic implications. Science 369, eabc8511 (2020).

4. M. Koutsakos et al., Integrated immune dynamics define correlates of COVID-19 severity and antibody responses. Cell Rep Med 2, 100208 (2021).

5. P. S. Arunachalam et al., Systems biological assessment of immunity to mild versus severe COVID-19 infection in humans. Science 369, 1210–1220 (2020).

6. L. Rodriguez et al., Systems-Level Immunomonitoring from Acute to Recovery Phase of Severe COVID-19. Cell Rep Med 1, 100078 (2020).

7. R. C. Group et al., Dexamethasone in Hospitalized Patients with Covid-19. N Engl J Med 384, 693–704 (2021).

8. A. C. Kalil et al., Baricitinib plus Remdesivir for Hospitalized Adults with Covid-19. N Engl J Med 384, 795–807 (2021).

9. R. C. Group, Tocilizumab in patients admitted to hospital with COVID-19 (RECOVERY): a randomised, controlled, open-label, platform trial. Lancet 397, 1637–1645 (2021).

10. D. M. Weinreich et al., REGN-COV2, a Neutralizing Antibody Cocktail, in Outpatients with Covid-19. N Engl J Med 384, 238–251 (2021).

11. R. L. Gottlieb et al., Effect of Bamlanivimab as Monotherapy or in Combination With Etesevimab on Viral Load in Patients With Mild to Moderate COVID-19: A Randomized Clinical Trial. JAMA 325, 632–644 (2021).

12. F. P. Polack et al., Safety and Efficacy of the BNT162b2 mRNA Covid-19 Vaccine. N Engl J Med 383, 2603–2615 (2020).

13. N. Dagan et al., BNT162b2 mRNA Covid-19 Vaccine in a Nationwide Mass Vaccination Setting. N Engl J Med 384, 1412–1423 (2021).

14. L. R. Baden et al., Efficacy and Safety of the mRNA-1273 SARS-CoV-2 Vaccine. N Engl J Med 384, 403–416 (2021).

15. M. Voysey et al., Safety and efficacy of the ChAdOx1 nCoV-19 vaccine (AZD1222) against SARS-CoV-2: an interim analysis of four randomised controlled trials in Brazil, South Africa, and the UK. Lancet 397, 99–111 (2021).

16. J. Sadoff et al., Safety and Efficacy of Single-Dose Ad26.COV2.S Vaccine against Covid-19. N Engl J Med 384, 2187–2201 (2021).

17. N. Pardi, M. J. Hogan, F. W. Porter, D. Weissman, mRNA vaccines - a new era in vaccinology. Nat Rev Drug Discov 17, 261–279 (2018).

18. E. Bettini, M. Locci, SARS-CoV-2 mRNA Vaccines: Immunological Mechanism and Beyond. Vaccines (Basel) 9, 147 (2021).

19. M. Jeyanathan et al., Immunological considerations for COVID-19 vaccine strategies. Nat Rev Immunol 20, 615–632 (2020).

20. P. S. Arunachalam et al., Systems vaccinology of the BNT162b2 mRNA vaccine in humans. Nature, doi: 10.1038/s41586-41021-03791-x (2021).

21. B. A. Woldemeskel, C. C. Garliss, J. N. Blankson, SARS-CoV-2 mRNA vaccines induce broad CD4+ T cell responses that recognize SARS-CoV-2 variants and HCoV-NL63. J Clin Invest 131, e149335 (2021).

22. A. Sattler et al., Impaired Humoral and Cellular Immunity after SARS-CoV2 BNT162b2 (Tozinameran) Prime-Boost Vaccination in Kidney Transplant Recipients. medRxiv, 2021.2004.2006.21254963 (2021).

23. U. Sahin et al., COVID-19 vaccine BNT162b1 elicits human antibody and TH1 T cell responses. Nature 586, 594–599 (2020).

24. S. M. Barone et al., Unsupervised machine learning reveals key immune cell subsets in COVID-19, rhinovirus infection, and cancer therapy. bioRxiv, DOI: 10.1101/2020.1107.1131.190454 (2020).

25. I. Setliff et al., High-Throughput Mapping of B Cell Receptor Sequences to Antigen Specificity. Cell 179, 1636–1646 e1615 (2019).

26. K. E. Diggins, A. R. Greenplate, N. Leelatian, C. E. Wogsland, J. M. Irish, Characterizing cell subsets using marker enrichment modeling. Nat Methods 14, 275–278 (2017).

27. A. T. Huang et al., A systematic review of antibody mediated immunity to coronaviruses: kinetics, correlates of protection, and association with severity. Nat Commun 11, 4704 (2020).

28. J. B. Case et al., Neutralizing Antibody and Soluble ACE2 Inhibition of a Replication-Competent VSV-SARS-CoV-2 and a Clinical Isolate of SARS-CoV-2. Cell Host Microbe 28, 475–485 e475 (2020).

29. P. Gilchuk et al., Standardized Two-Step Testing of Antibody Activity in COVID-19 Convalescent Plasma. Cell Rep Med https://ssrn.com/abstract=3878407 DOI: 10.2139/ssrn.3878407 (2021).

30. S. J. Zost et al., Potently neutralizing and protective human antibodies against SARS-CoV-2. Nature 584, 443–449 (2020).

31. D. F. Robbiani et al., Convergent antibody responses to SARS-CoV-2 in convalescent individuals. Nature 584, 437–442 (2020).

32. T. F. Rogers et al., Isolation of potent SARS-CoV-2 neutralizing antibodies and protection from disease in a small animal model. Science 369, 956–963 (2020).

33. Z. Wang et al., mRNA vaccine-elicited antibodies to SARS-CoV-2 and circulating variants. Nature 592, 616–622 (2021).

34. A. R. Shiakolas et al., Cross-reactive coronavirus antibodies with diverse epitope specificities and Fc effector functions. Cell Rep Med 2, 100313 (2021).

35. M. I. J. Raybould, A. Kovaltsuk, C. Marks, C. M. Deane, CoV-AbDab: the coronavirus antibody database. Bioinformatics 37, 734–735 (2021).

36. E. C. Chen et al., Convergent antibody responses to the SARS-CoV-2 spike protein in convalescent and vaccinated individuals. bioRxiv, DOI:10.1101/2021.1105.1102.442326 (2021).

37. M. I. Samanovic et al., Poor antigen-specific responses to the second BNT162b2 mRNA vaccine dose in SARS-CoV-2-experienced individuals. medRxiv, DOI:10.1101/2021.1102.1107.21251311 (2021).

38. R. S. Herati et al., Successive annual influenza vaccination induces a recurrent oligoclonotypic memory response in circulating T follicular helper cells. Sci Immunol 2, eaag2152 (2017).

39. A. Cardeno, M. K. Magnusson, M. Quiding-Jarbrink, A. Lundgren, Activated T follicular helper-like cells are released into blood after oral vaccination and correlate with vaccine specific mucosal B-cell memory. Sci Rep 8, 2729 (2018).

40. J. E. Huber et al., Dynamic changes in circulating T follicular helper cell composition predict neutralising antibody responses after yellow fever vaccination. Clin Transl Immunology 9, e1129 (2020).

41. L. M. Muehling et al., Single-Cell Tracking Reveals a Role for Pre-Existing CCR5+ Memory Th1 Cells in the Control of Rhinovirus-A39 After Experimental Challenge in Humans. J Infect Dis 217, 381–392 (2018).

42. Z. C. Wang et al., Extrafollicular PD-1(high)CXCR5(-)CD4(+) T cells participate in local immunoglobulin production in nasal polyps. J Allergy Clin Immunol S0091-6749, 01050–01052 (2021).

43. J. S. Hale, R. Ahmed, Memory T follicular helper CD4 T cells. Front Immunol 6, 16 (2015).

44. X. Zhu, J. Zhu, CD4 T Helper Cell Subsets and Related Human Immunological Disorders. Int J Mol Sci 21, 8011 (2020).

45. D. Fang et al., Transient T-bet expression functionally specifies a distinct T follicular helper subset. J Exp Med 215, 2705–2714 (2018).

46. D. S. Khoury et al., Neutralizing antibody levels are highly predictive of immune protection from symptomatic SARS-CoV-2 infection. Nat Med 27, 1205–1211 (2021).

47. A. J. McMichael, F. M. Gotch, G. R. Noble, P. A. Beare, Cytotoxic T-cell immunity to influenza. N Engl J Med 309, 13–17 (1983).

48. A. B. Rickinson, D. J. Moss, Human cytotoxic T lymphocyte responses to Epstein-Barr virus infection. Annu Rev Immunol 15, 405–431 (1997).

49. J. R. Currier et al., A panel of MHC class I restricted viral peptides for use as a quality control for vaccine trial ELISPOT assays. J Immunol Methods 260, 157–172 (2002).

50. L. D. Estrada, S. Schultz-Cherry, Development of a Universal Influenza Vaccine. J Immunol 202, 392–398 (2019).

51. A. J. McMichael, Is a Human CD8 T-Cell Vaccine Possible, and if So, What Would It Take? Could a CD8(+) T-Cell Vaccine Prevent Persistent HIV Infection? Cold Spring Harb Perspect Biol 10, a029124 (2018).

52. M. D. Co, E. D. Kilpatrick, A. L. Rothman, Dynamics of the CD8 T-cell response following yellow fever virus 17D immunization. Immunology 128, e718–727 (2009).

53. S. A. Fuertes Marraco et al., Long-lasting stem cell-like memory CD8+ T cells with a naive-like profile upon yellow fever vaccination. Sci Transl Med 7, 282ra248 (2015).

54. A. H. Ellebedy et al., Defining antigen-specific plasmablast and memory B cell subsets in human blood after viral infection or vaccination. Nat Immunol 17, 1226–1234 (2016).

55. G. E. Hartley et al., Rapid generation of durable B cell memory to SARS-CoV-2 spike and nucleocapsid proteins in COVID-19 and convalescence. Sci Immunol 5, eabf8891 (2020).

56. F. Ahmadizar et al., QTc-interval prolongation and increased risk of sudden cardiac death associated with hydroxychloroquine. Eur J Prev Cardiol, zwaa118 (2020).

57. E. M. Anderson et al., Seasonal human coronavirus antibodies are boosted upon SARS-CoV-2 infection but not associated with protection. Cell 184, 1858–1864 e1810 (2021).

58. P. Kaplonek et al., Early cross-coronavirus reactive signatures of protective humoral immunity against COVID-19. bioRxiv, DOI:10.1101/2021.1105.1111.443609 (2021).

59. A. R. Parker, M. A. Park, S. Harding, R. S. Abraham, The total IgM, IgA and IgG antibody responses to pneumococcal polysaccharide vaccination (Pneumovax(R)23) in a healthy adult population and patients diagnosed with primary immunodeficiencies. Vaccine 37, 1350–1355 (2019).

60. R. J. Cox, K. A. Brokstad, P. Ogra, Influenza virus: immunity and vaccination strategies. Comparison of the immune response to inactivated and live, attenuated influenza vaccines. Scand J Immunol 59, 1–15 (2004).

61. B. A. Halperin et al., Kinetics of the antibody response to tetanus-diphtheria-acellular pertussis vaccine in women of childbearing age and postpartum women. Clin Infect Dis 53, 885–892 (2011).

62. D. Sterlin et al., IgA dominates the early neutralizing antibody response to SARS-CoV-2. Sci Transl Med 13, eabd2223 (2021).

63. A. V. Wisnewski, J. Campillo Luna, C. A. Redlich, Human IgG and IgA responses to COVID-19 mRNA vaccines. PLoS One 16, e0249499 (2021).

64. F. Meiler, S. Klunker, M. Zimmermann, C. A. Akdis, M. Akdis, Distinct regulation of IgE, IgG4 and IgA by T regulatory cells and toll-like receptors. Allergy 63, 1455–1463 (2008).

65. N. Zheng et al., TLR7 in B cells promotes renal inflammation and Gd-IgA1 synthesis in IgA nephropathy. JCI Insight 5, e136965 (2020).

66. J. Bessa et al., Alveolar macrophages and lung dendritic cells sense RNA and drive mucosal IgA responses. J Immunol 183, 3788–3799 (2009).

67. Y. H. Kim et al., Inactivated influenza vaccine formulated with single-stranded RNA-based adjuvant confers mucosal immunity and cross-protection against influenza virus infection. Vaccine 38, 6141–6152 (2020).

68. Y. Peled et al., BNT162b2 vaccination in heart transplant recipients: Clinical experience and antibody response. J Heart Lung Transplant S1053-2498, 02274–02279 (2021).

69. A. Sattler et al., Impaired humoral and cellular immunity after SARS-CoV-2 BNT162b2 (tozinameran) prime-boost vaccination in kidney transplant recipients. J Clin Invest 131, e150175 (2021).

70. S. Marinaki et al., Immunogenicity of SARS-CoV-2 BNT162b2 vaccine in solid organ transplant recipients. Am J Transplant, DOI: 10.1111/ajt.16607 (2021).

71. P. Grivas et al., Association of clinical factors and recent anticancer therapy with COVID-19 severity among patients with cancer: a report from the COVID-19 and Cancer Consortium. Ann Oncol 32, 787–800 (2021).

72. V. Furer et al., Immunogenicity and safety of the BNT162b2 mRNA COVID-19 vaccine in adult patients with autoimmune inflammatory rheumatic diseases and in the general population: a multicentre study. Ann Rheum Dis, DOI: 10.1136/annrheumdis-2021-220647 (2021).

73. S. K. Mahil et al., The effect of methotrexate and targeted immunosuppression on humoral and cellular immune responses to the COVID-19 vaccine BNT162b2: a cohort study. Lancet Rheumatol, DOI: 10.1016/S2665-9913(1021)00212-00215 (2021).

74. D. H. Huson, C. Scornavacca, Dendroscope 3: an interactive tool for rooted phylogenetic trees and networks. Syst Biol 61, 1061–1067 (2012).

## Methods References

1. P. B. Ferrell, Jr. et al., High-Dimensional Analysis of Acute Myeloid Leukemia Reveals Phenotypic Changes in Persistent Cells during Induction Therapy. PLoS One 11, e0153207 (2016).

2. A. R. Greenplate et al., Myelodysplastic Syndrome Revealed by Systems Immunology in a Melanoma Patient Undergoing Anti-PD-1 Therapy. Cancer Immunol Res 4, 474–480 (2016).

3. R. Finck et al., Normalization of mass cytometry data with bead standards. Cytometry Part A 83A, 483–494 (2013).

4. J. M. Irish et al., B-cell signaling networks reveal a negative prognostic human lymphoma cell subset that emerges during tumor progression. Proceedings of the National Academy of Sciences 107, 12747 (2010).

5. C. E. Roe, M. J. Hayes, S. M. Barone, J. M. Irish, Training Novices in Generation and Analysis of High-Dimensional Human Cell Phospho-Flow Cytometry Data. Current Protocols in Cytometry 93, e71 (2020).

6. S. M. Barone et al., Unsupervised machine learning reveals key immune cell subsets in COVID-19, rhinovirus infection, and cancer therapy. bioRxiv, 2020.2007.2031.190454 (2020).

7. K. E. Diggins, A. R. Greenplate, N. Leelatian, C. E. Wogsland, J. M. Irish, Characterizing cell subsets using marker enrichment modeling. Nat Methods 14, 275–278 (2017).

8. J. R. Cantor et al., Physiologic Medium Rewires Cellular Metabolism and Reveals Uric Acid as an Endogenous Inhibitor of UMP Synthase. Cell 169, 258–272 e217 (2017).

9. I. Setliff et al., High-Throughput Mapping of B Cell Receptor Sequences to Antigen Specificity. Cell 179, 1636–1646.e1615 (2019).

10. A. R. Shiakolas et al., Cross-reactive coronavirus antibodies with diverse epitope specificities and Fc effector functions. Cell Rep Med 2, 100313 (2021).

11. E. Alamyar, P. Duroux, M. P. Lefranc, V. Giudicelli, IMGT((R)) tools for the nucleotide analysis of immunoglobulin (IG) and T cell receptor (TR) V-(D)-J repertoires, polymorphisms, and IG mutations: IMGT/V-QUEST and IMGT/HighV-QUEST for NGS. Methods Mol Biol 882, 569–604 (2012).

12. N. T. Gupta et al., Change-O: a toolkit for analyzing large-scale B cell immunoglobulin repertoire sequencing data. Bioinformatics 31, 3356–3358 (2015).

13. D. H. Huson, C. Scornavacca, Dendroscope 3: an interactive tool for rooted phylogenetic trees and networks. Syst Biol 61, 1061–1067 (2012).

14. R. Satija, J. A. Farrell, D. Gennert, A. F. Schier, A. Regev, Spatial reconstruction of single-cell gene expression data. Nat Biotechnol 33, 495–502 (2015).

